# Single nucleus RNA-seq in the hippocampus of a Down syndrome mouse model reveals new key players in memory

**DOI:** 10.1101/2021.11.18.469102

**Authors:** Cesar Sierra, Ilario De Toma, Mara Dierssen

**Affiliations:** Center for Genomic Regulation, The Barcelona Institute for Science and Technology, Barcelona, Spain; Neurosciences Research Program, Hospital del Mar Medical Research Institute, Barcelona, Spain; University Pompeu Fabra, Barcelona, Spain; Centro de Investigación Biomédica en Red de Enfermedades Raras, Barcelona, Spain

## Abstract

Down syndrome (DS) is the most common genetic cause of intellectual disability. Even though great advances in the last decades have allowed better delineation of its pathogenetic mechanisms, its cellular and molecular bases are still poorly understood. To evaluate the consequences of chromosome aneuploidy on the hippocampus, we analyzed single-nucleus transcriptional profiles of the DS mouse model Ts65Dn. Our data revealed abnormal cell composition specifically of the Ts65Dn dentate gyrus and of specific subtypes of interneurons, without major changes in CA1 or CA3. We found that trisomy results in a highly cell-type specific global alteration of the transcriptome and detected previously undefined differentially expressed genes in specific neuronal populations. We identified the long-non-coding gene Snhg11 to be specifically downregulated in the trisomic dentate gyrus and provide evidence for its involvement in hippocampal-dependent cognitive phenotypes, possibly contributed by impaired adult neurogenesis.

## Introduction

Down syndrome (DS) is caused by trisomy of Homo sapiens chromosome 21 (HSA21) and is the most common cause of genetic intellectual disability, affecting more than 5 million people globally. DS alters central nervous system development, impairing cognition, and adaptive behavior. It has an extremely heterogeneous and complex pathophysiology, with a number of cellular and molecular pathways contributing to the learning and memory impairment. Neuropsychological investigations indicate that deficits in hippocampal-mediated learning and memory processes are hallmarks of DS [1, 2], and molecular and cellular defects have been detected in post-mortem fetal DS hippocampus [3, 4].

Mouse models of DS, carrying various lengths of chromosomal duplications, also recapitulate most of the DS hippocampal neuropathology [4, 5]. In particular, models carrying some of the largest duplications, Ts65Dn (TS) and Dp(16)1Yey, display hippocampal learning and memory deficits associated with reduced hippocampal long-term potentiation [6-9], and alterations in the organization of spiking of hippocampal CA1 pyramidal neurons [10] thus mimicking the human condition in regards to hippocampal dependent deficits. Other mechanisms, such as altered neurogenesis have also been proposed to be neurobiological correlates of intellectual disability in DS. Cell proliferation from early postnatal stages is reduced in the subgranular zone of fetuses with DS and of Ts65Dn mice [3, 11]. The number of differentiated neurons is also reduced in individuals with DS [12], although the number of astrocytes remains unaltered [13]. This impairment continues into adulthood and may participate in learning and memory deficits ([14, 15].

Besides, an increasing number of studies suggest that an excitatory-inhibitory imbalance occurs in DS [16], potentially contributing to synaptic and circuit disorganization in DS and is also implicated in DS learning deficits. This imbalance might be in part explained by the increased number of certain subtypes of GABAergic interneurons in the trisomic hippocampus [17, 18]. However, comprehensive assessments of changes in cellular composition of the DS brain have not been possible to date, owing to limitations of the available assays.

DS is a disorder of gene expression deregulation, as the triplication of HSA21 results in a global disturbance of the transcriptome that is proposed to contribute to the phenotypic manifestations of DS [19]. This global gene expression deregulation is likely caused by alterations intrinsic to the extra copy of HSA21, such as the overexpression of genes involved in epigenetic regulation. This is the case of *DNMT3L, MIS18A, N2AMT1, DYRK1A, BRWD1, RUNX1, HLCS, HMGN1* and *ETS2*. In fact, several studies have suggested chromatin dysfunction in DS [20-22]. However, other possible mechanisms associated with the regulation of chromatin function are still unexplored. For example, despite HSA21 being the smallest autosome, it is highly enriched in long-non coding RNAs (lncRNAs) [23], and a high number of lncRNAs are abnormally expressed in DS induced pluripotent stem cells (iPSCs) [24]. Interestingly, lncRNAs have been associated with learning and memory [25-27] and adult neurogenesis [28-30], but little is known about their direct mechanisms. While some studies suggest their role in the proliferation of neural progenitor cells [30], other work proposes that lncRNAs modulate their ability to differentiate [29], or the migration of newly born cells to their proper location within the brain [29]. Interestingly, lncRNAs might also be critical for the adult hippocampal GABAergic circuit [31], thus covering several of the DS pathogenicity mechanisms. While lncRNAs are generally detected at low levels in bulk tissues, single-cell transcriptomics revealed that lncRNAs are abundantly expressed in individual cells and are highly cell type-specific [32].

Here, in order to characterize the cellular and molecular alterations associated with DS hippocampal dysfunction, we explored the cellular composition of the hippocampus of the DS mouse model Ts65Dn, using single nucleus RNA sequencing to dissect neural cell diversity as well as dysregulation of molecular signatures associated with specific cell types. Our study revealed a highly cell-type specific global alteration of the transcriptome and detected previously unknown differentially expressed genes in specific neuronal populations. Strikingly, we identified *Snhg11* to be specifically downregulated in the trisomic dentate gyrus and provide evidence for its involvement in hippocampal-dependent memory and adult neurogenesis. To our knowledge, this is the first single nuclei sequencing study on a trisomic hippocampus, characterizing the transcriptome of tens of thousands of individual hippocampal neurons in parallel and thereby offering a cell and molecular atlas of the trisomic hippocampus.

## Results

### Unbiased identification of neuronal subtypes in hippocampus

To address the question of how the different neuronal populations are affected in the TS hippocampus, we isolated the NeuN+ population by fluorescence activated nuclear sorting (FANS; see Methods). After FANS, using 10X we sequenced 27602 and 28545 nuclei with high integrity from four WT and four TS animals, respectively (Fig. 1A). Since single nuclei RNA-seq (snRNA-seq) profiles nuclear RNA, our gene expression profile data reflect nascent transcription, as well as the cellular transcriptome.

**Figure 1.**
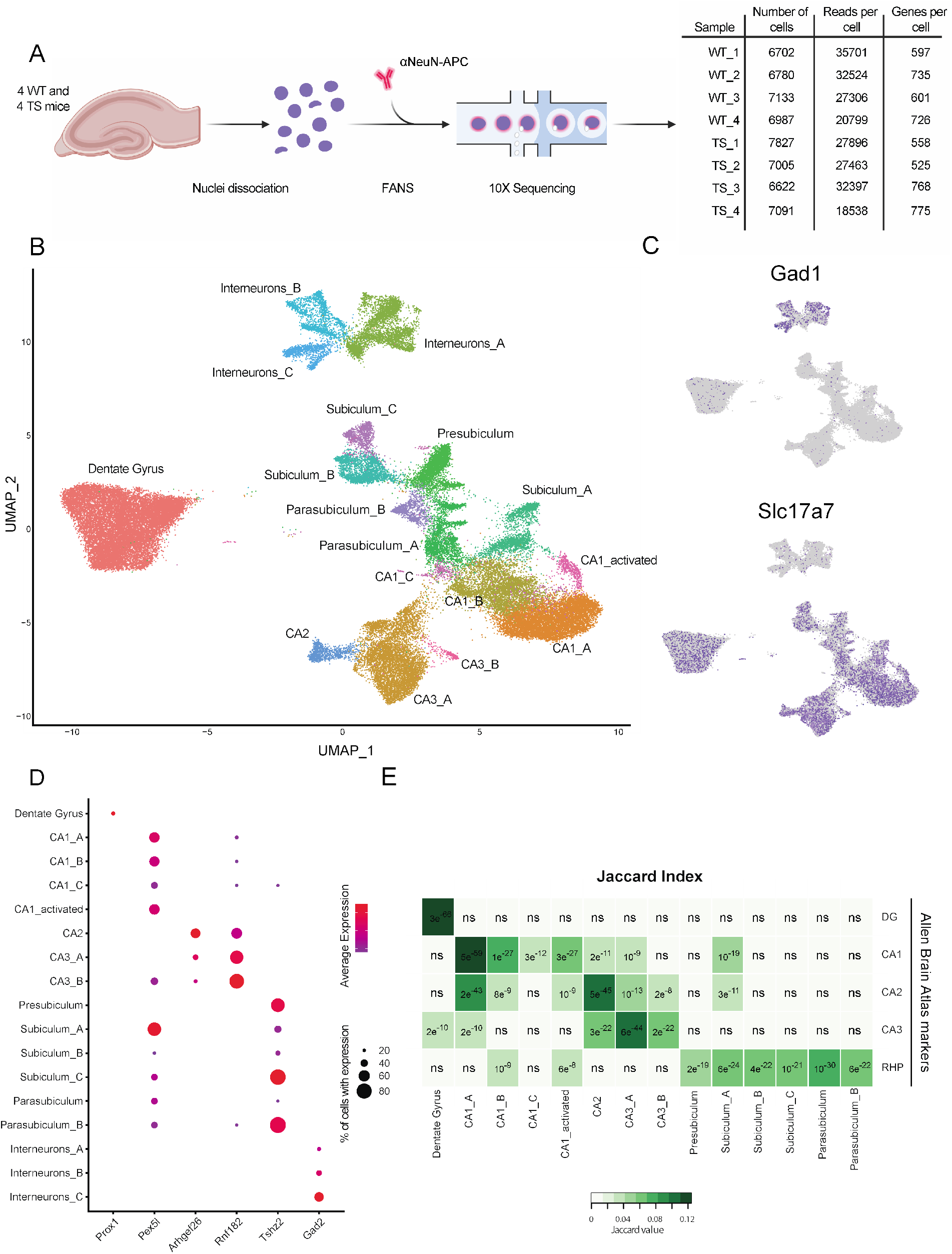
Unbiased identification of neuronal subtypes in hippocampus. **(A)** Overview of the experimental strategy with schematic depicting isolation strategy for NeuN-positive cells. **(B)** UMAP embedding of all hippocampal single nuclei showing cell clusters. Cells are colored by cell subtype. **(C)** Mapping of known cell markers of excitatory (*Slc17a7*) and inhibitory (*Gad1*) neurons. **(D)** Dotplot showing enrichment of region-specific markers in each cluster. **(E)** Jaccard overlap between cluster markers and genes enriched in each hippocampal subregion according to Allen Brain Atlas. p-values are indicated.

Single-cell feature-barcode matrices were used to embed cells in a K-nearest neighbor graph that defines cell clusters in an unbiased manner. Nuclear transcriptomes were visualized using a uniform manifold approximation and projection (UMAP) plot (Fig. 1B). We detected 17 clusters of cells sharing similar gene expression patterns. These clusters did not result from technical or batch effects (Supplementary Fig. 1).

To determine the identity of each cluster, we obtained the gene markers of each cluster based on their differential expression with respect to the remaining clusters. Overall, we identified a total of 1191 marker genes for the different hippocampal neuron subpopulations (Supplementary Table 1). The comparison of these markers to known signatures of hippocampal neuronal types allowed the identification of the excitatory and interneuron populations. Classical gene markers for these populations, such as *Slc17a7* for excitatory neurons, *Gad1* for interneurons showed a clear enrichment in specific clusters (Fig. 1C).

Glutamatergic cells were further mapped to hippocampal subregions, namely Dentate Gyrus (DG), CA1, CA2, CA3 and Retrohippocampal region (RHP). We defined the identities of each neuronal subtypes by analyzing the subregional expression pattern of their top marker genes (Fig. 1D) in the Allen Brain Atlas [33] (Supplementary Fig. 2A). Thereby, we identified *Prox1* as a marker of DG, *Pex5l* of CA1, *Arhgef26* of CA2, *Rnf182* of CA3, and *Tshz* of RHP. To validate this mapping in a more systematic manner, we studied the overlap between the top 300 marker genes of each of our clusters and the 300 most differentially expressed genes in each hippocampal subregion identified by the Allen Brain Atlas [33] (Fig. 1E). Using this same approach, we were also able to identify the major subregions of the RHP, namely presubiculum, subiculum and parasubiculum (Supplementary Fig. 2B).

### Granule cells and specific interneurons subtypes are altered in the trisomic hippocampus

The compositional analysis via bootstrap [34] of the main subregions in the WT and TS replicates revealed a neuron subtype-specific alteration of the cellular composition of the TS hippocampus. Specifically, the granule cell population from the DG was reduced in the trisomic hippocampus compared to WT. Instead, the overall interneuron population was increased in trisomics (Fig. 2A, Supplementary Fig. 3A). Given that over-inhibition leads to cognitive deficits in TS mice [4, 16, 35, 36], possibly due to overproduction of specific interneuron subtypes [17, 18], we further investigated how the trisomy may impact specific subclusters of interneurons. To this aim we subsetted this population and increased the resolution in cluster identification. This approach resulted in a total of five inhibitory subclusters (Fig. 2B). To identify them, we overlapped their top marker genes (Supplementary Table 2) with the marker genes of the most represented hippocampal interneuron clusters in mouse brain [37] (Fig. 2C).

**Figure 2.**
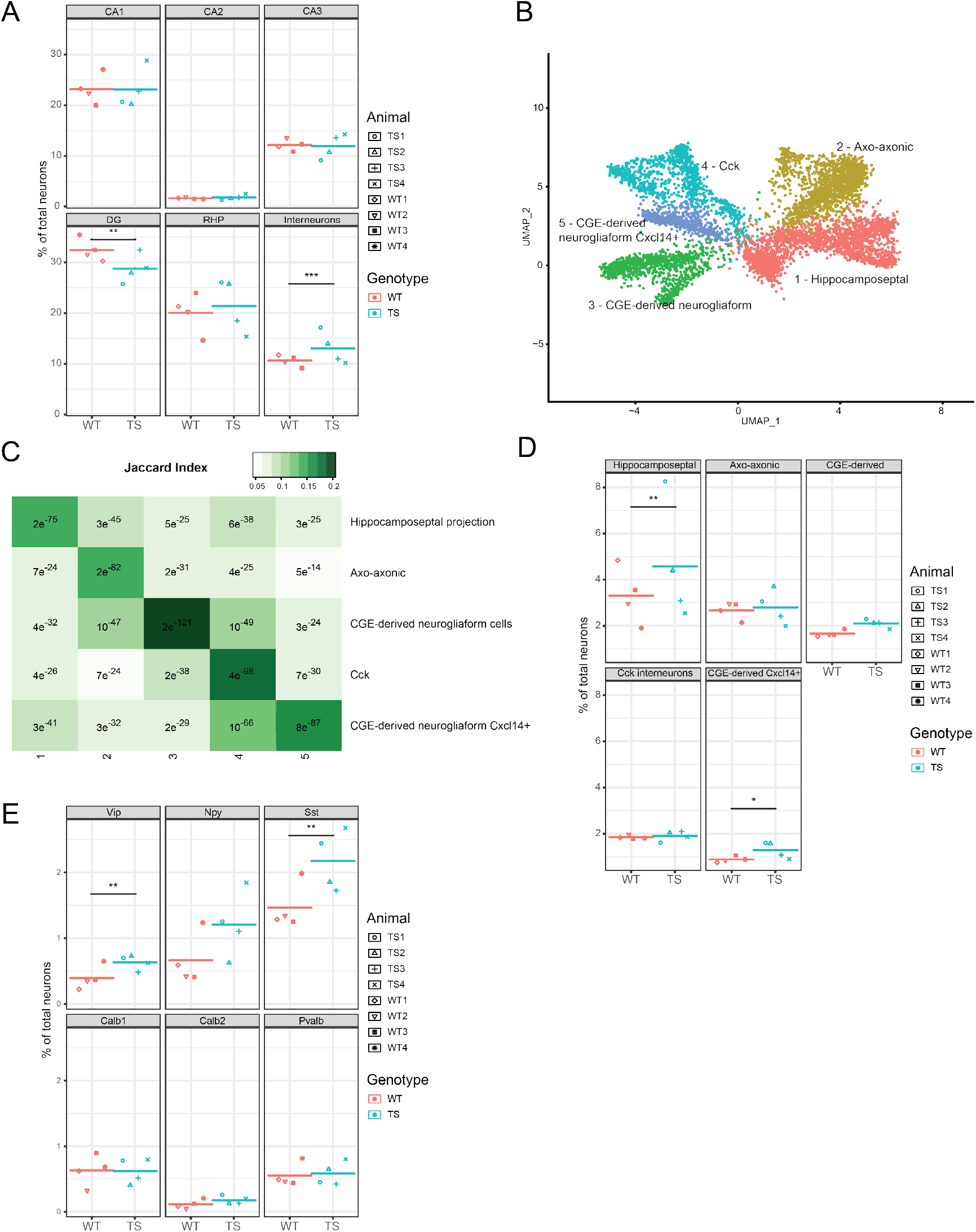
Compositional analysis of hippocampal subpopulation in WT and TS hippocampus. **(A)** Dotplot showing the percentage of WT (red) and TS (blue) cells belonging to the main neuronal subtypes. Each symbol represents an individual (TS1-4; WT1-4). ** p < 0.01, *** p < 0.001; Generalized linear model. RHP: retrohippocampal neurons. **(B)** UMAP plot showing subsetted interneurons at an increased resolution. CGE: caudal ganglionic eminence. **(C)** Jaccard overlaps between cluster markers of the dataset and the markers of the five major hippocampal interneuron populations described in Zeiset *et al*. [37]. **(D)** Dotplot showing the percentage of WT(red) and TS (blue) cells belonging to each of the five interneurons subclusters identified. * p < 0.05, ** p < 0.01; Generalized linear model **(E)** Dotplot of the percentage of WT and TS cells expressing the major classical interneuron markers. ** p < 0.01; Student t-test.

Cluster 1 was identified with Sst+ hippocamposeptal interneurons, expressing *Sst* and *Grm1* [38]. Cluster 2 was identified as axo-axonic cells, also known as chandelier cells, because of their high expression of *Pvalb*. However, this cluster probably contains a subpopulation of basket cells, given the presence of markers such as *Satb1* and *Erbb4* and the lack of expression of Grm1. Cluster 3 showed a clear identity as caudal ganglionic eminence (CGE)-derived neurogliaform cells as determined by the high expression of *Cacna2d1* along with low expression of *Lhx6*. Cluster 4 could be identified as *Cck* interneurons, probably located outside of the stratum radiatum / stratum lacunosum-moleculare given the lack of expression of *Cxcl14*. Lastly, the smallest interneuron cluster did not have any top marker that allowed its direct identification. However, this cluster showed a very significant overlap with the top markers of CGE-derived neurogliaform *Cxcl14*+ cells in the adolescent mouse brain dataset [37].

The quantification of the cluster sizes of the four WT and four trisomic animals showed subtype-specific changes in interneuron abundance (Fig. 2D, Supplementary Fig. 3B). Interestingly, the increase in the inhibitory population was found to be specific to hippocamposeptal interneurons and

CGE-derived neurogliaform *Cxcl14*+ cells. Instead, axo-axonic and *Cck* interneurons showed similar abundances in WT and TS hippocampus, and CGE-derived neurogliaform cells presented a non-significant increase in TS.

We also quantified the fraction of interneurons expressing each of the neuropeptides that characterize the different interneuron subtypes. This led to the identification of *Vip*+ and *Sst*+ cells as the only interneuron subtypes significantly increased in the trisomic hippocampus (Fig. 2E) while other markers such as *Calb1, Calb2, Pvalb* and *Npy* are unaltered.

### The transcriptome alteration induced by the trisomy is highly cell-type specific

The differential expression analyses between the trisomic and euploid major neuronal subtypes (CA1, CA2, CA3, DG, RHP and interneurons) showed a total of 293 differentially expressed genes (DEG; Fig. 3A, Supplementary Table 3). As expected, the highest portion of DEGs, mostly upregulated, were located to the mouse chromosome 16 (Mmu16) (Fig. 3B), a portion of which is triplicated in Ts65Dn. We found that almost half (48%) of the DEGs significantly altered by the trisomy were cell subtype specific (Fig. 3C). Among DEG found in more than one neuronal subtype, the highest number of shared DEGs was found between CA1 and CA3, reflecting their close biological identity and functionality. As expected, a decreasing number of genes was found to be commonly deregulated in a higher number of cell subtypes, being the triplicated genes more prone to be commonly deregulated (Supplementary Fig. 4A). The high cell-specific impact of the trisomy is also illustrated in Figure 3D, showing that most of the DEGs identified at the single nuclei level were masked in the pseudobulk analysis of the same samples.

**Figure 3.**
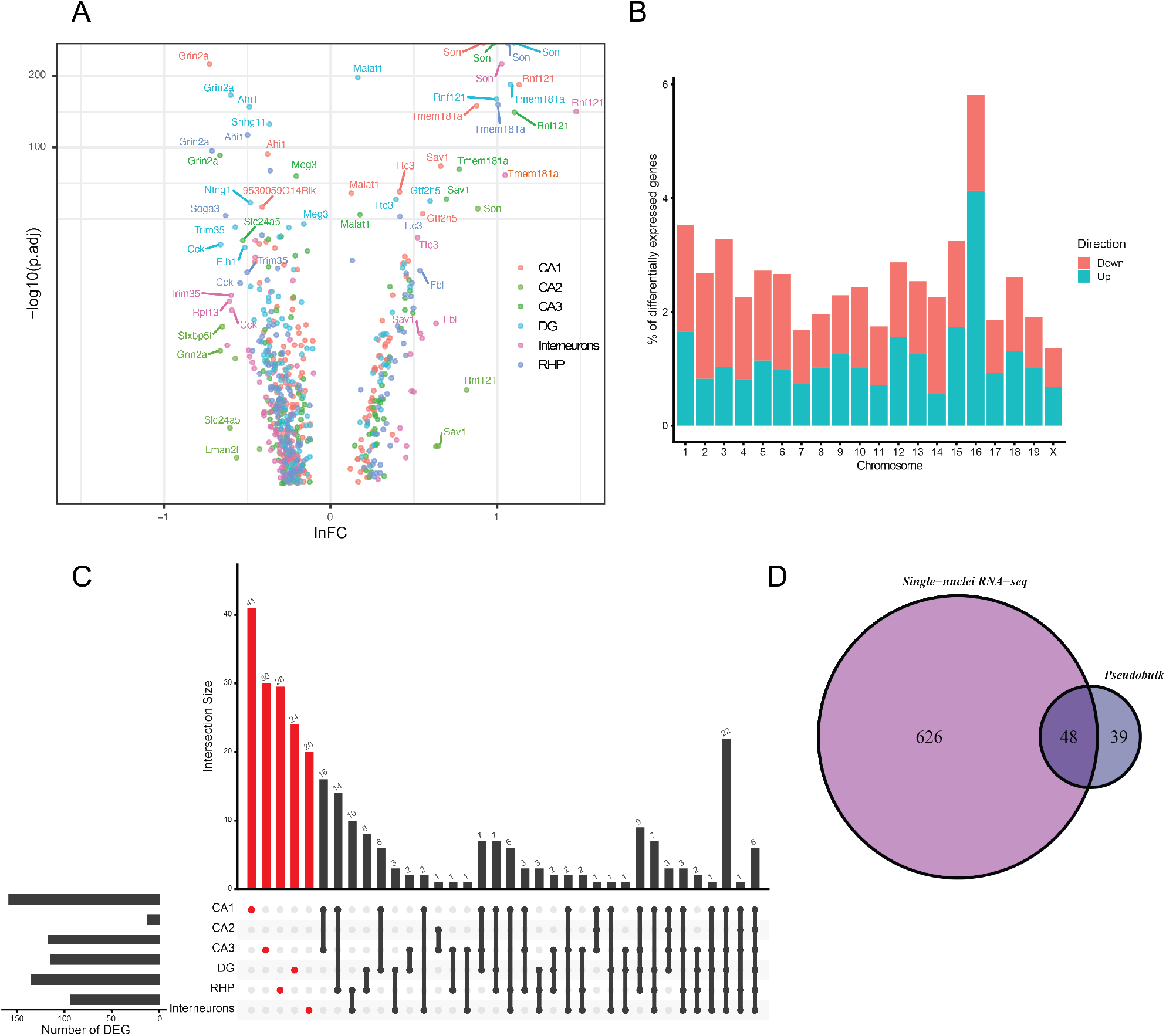
Differentially expressed genes (DEGs) induced by trisomy in hippocampal cell types. **(A)** Volcano Plot of DEGs colored by each major neuronal subtype. Shown are names of DEGs with a lnFC < 0.4 or -log10(p.adj) > 60. **(B)** Distribution of DEGs along the mouse chromosomes. **(C)** DEGs unique to a cell type/subtype are indicated in red and those shared between 2 or more cell types or subtypes are indicated by black dots. The histogram above each plot indicates the number of DEGs for each cell type or subtype. The barplots on the left show the total number of DEGs per cellular type and subtype. **(D)** Many cell-type and subtype specific DEGs are not captured in pseudobulk analysis.

The identification of cell type-specific DEGs resulting from the trisomy might be of interest to elucidate the mechanisms that lead to DS hippocampal dysfunction (Table 1 and Supplementary Fig. 4B). For instance, Epha6, a key regulator of neuronal and spine morphology, was specifically downregulated in CA1 trisomic neurons. Instead, *Eid1*, which has been associated with an impaired synaptic plasticity of CA1 pyramidal neurons, is upregulated in this region [39]. Interestingly, we specifically found altered in the DG genes that are related to neurogenesis (Table 1, Supplementary Fig. 4C), which is severely affected in DS [14, 40, 41]. This is the case of the overexpression of *Flrt3*, which is associated with neuronal migration inhibition [42] and the downregulation of *Ptprd*, which is a critical regulator of neurogenesis [43]. The enrichment of genes related to adult neurogenesis among DEGs of the trisomic DG might explain the deficits observed in this process in DS. Two voltage-gated K+ channel subunits, Kv1.1 and Kv3.1, encoded by *Kcna1* and *Kcnc1*, respectively, were specifically downregulated in interneurons. This reduced expression could lead to an hyperexcitability of trisomic interneurons, thereby contributing to the imbalance in excitatory/inhibitory transmission in the trisomic brain.

**Table 1.**
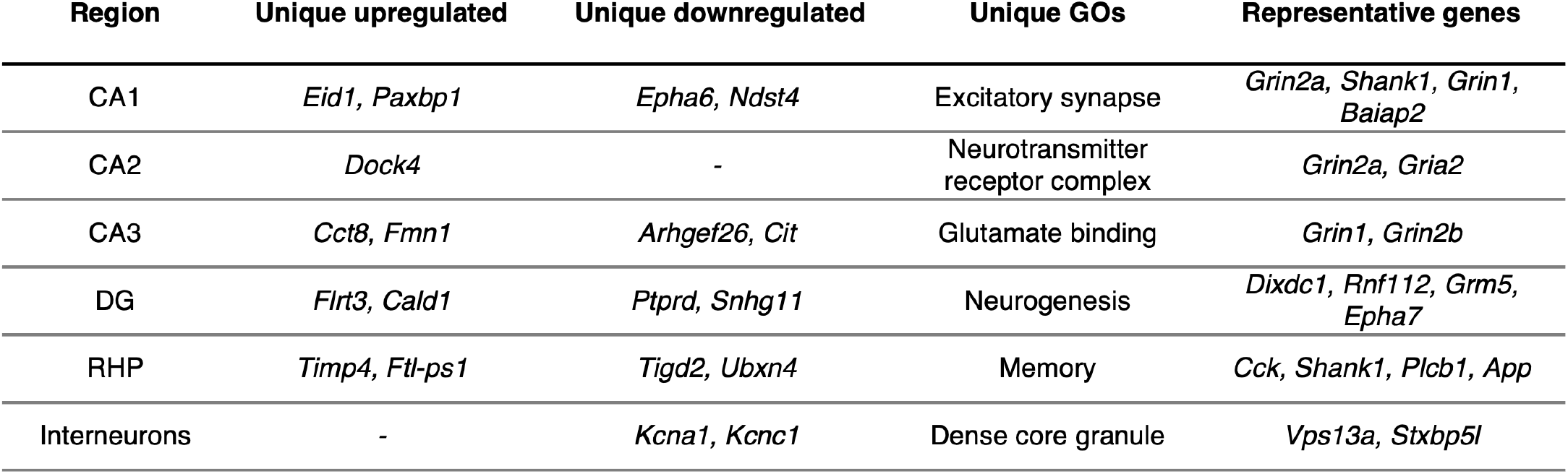
Top up- and downregulated genes and top enriched Gene Ontologies unique for each major neuronal subtype.

Not all triplicated genes were equally altered in each cellular subtype. Whereas *Gart* is uniquely overexpressed in trisomic CA3 neurons, Son is consistently greatly overexpressed in every cellular type examined. Among downregulated genes, we found *Grin2a*, encoding the Glutamate Ionotropic Receptor NMDA Type Subunit 2A, to be dramatically reduced in the whole trisomic hippocampus regardless of cellular types.

### The trisomy specifically impacts the DG transcriptomic profile

To visualize cell-subtype specific changes in the transcriptomic profiles of WT and trisomic neurons, we overlapped the UMAP plot from each genotype (Fig 4A). Strikingly, we observed a major shift in the two dimensional embedding of trisomic granule cells from the DG compared to WT, whereas the other neuronal subtypes overlapped between genotypes. This shift was quantified by testing if the euclidean distance of gene expression profiles of cells between the two genotypes is larger than expected by chance, confirming the observed shift in the DG and revealing a milder shift in CA1 (Supplementary Fig. 5A).

**Figure 4.**
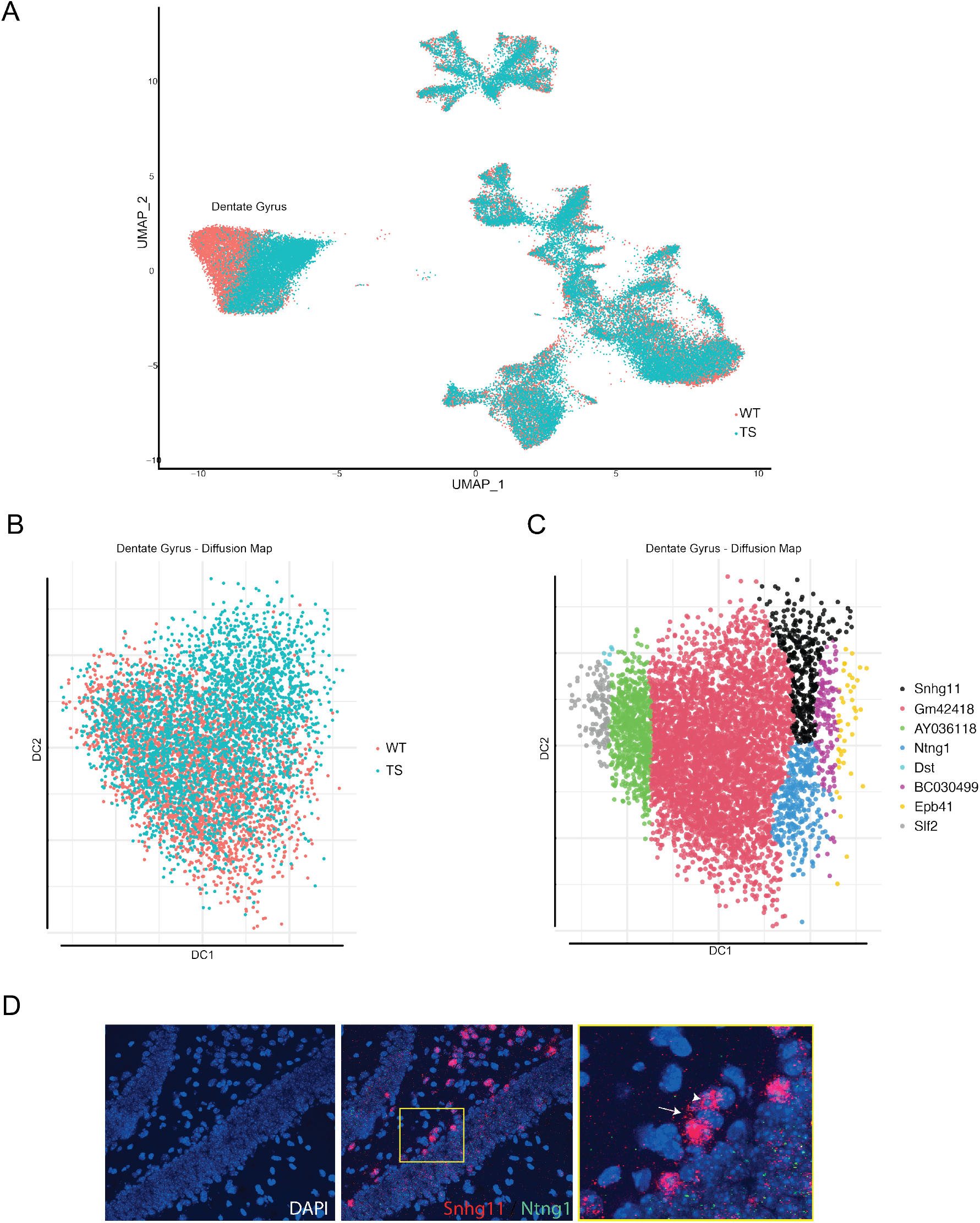
Transcriptomic shift specific to the dentate. **(A)** UMAP plot of WT (red) and trisomic (blue) cells. **(B)** Representation of the two-dimensional embedding of WT trisomic granule cells by Diffusion map. **(C)** Global and local gene relevance in the diffusion space. **(D)** Representative images of RNAscope in situ hybridization of Snhg11 and Ntng1 in the DG. Each individual 1-2μm dot, represents an individual RNA molecule and its location within cells. Arrow: cytoplasmic localization of Snhg11. Arrow head: nuclear localization of Snhg11.

To validate this transcriptomic shift, we subset the DG nuclei and repeated the clustering and the two-dimensional reduction by an independent approach, the Diffusion Map. Similarly to UMAP, we also observed a transcriptomic shift that is specific to the trisomic DG (Fig. 4B), while other subregions and cell types showed a high overlap between genotypes (Supplementary Fig. 5B). When further investigating this shift, we identified the small nucleolar RNA host gene 11, *Snhg11* as the gene with the highest global and local gene relevance in the DG diffusion map (Fig. 4C). This gene is a member of the non-protein-coding multiple snoRNA host gene family. Interestingly, this long non-coding gene is specifically differentially expressed in the trisomic DG, where its expression is strongly reduced (Table 1 and Fig. 3A). Similarly, Netrin G1, *Ntng1* is also dramatically downregulated in this subregion (Fig. 3A) and shows a high global gene relevance in the two dimensional embedding shift of the trisomic DG. *Ntng1* is a known marker of mature granule cells, which led us to think that cellular and functional identity of trisomic granule cells could be compromised. In fact, although the cluster-specific marker genes shared between genotypes showed a high overlap with the DG markers defined by the Allen Brain Atlas [33] (Fig. 1E), when we repeated this approach by genotype, we observed a significant loss of marker genes in the trisomic DG. This reduction of marker identity was specific to the DG, suggesting that the identity of granule cells is compromised (Supplementary Fig. 6).

The next question was about the possible role of Snhg11 in DS. In fact its expression and its role in the brain remain largely uncharacterized. To uncover its distribution in the DG, we used in situ hybridization to detect *Snhg11* and *Ntng1* in the WT DG. As previously reported, *Ntng1* signal appeared distributed along the granule cell layer as punctuated and with a cytoplasmic localization distributed along the granule cell layer (Fig. 4D). Similarly to most small nucleolar RNA host genes (SNHGs) [44], we found *Snhg11* in the nucleus as well as in the cytoplasm. Interestingly, although *Snhg11* signal is evenly distributed in the granule cell layer, certain cells localized to the subgranular layer display a particularly high expression of *Snhg11*, suggesting a potential role of this gene on adult neurogenesis.

### ASO-mediated knockdown of Snhg11 in vitro reduces cell proliferation

To investigate the function of *Snhg11* in the DG we knocked down its expression in the WT DG. To this aim, we designed two different antisense oligonucleotides (ASO) targeting the conserved exon 5 (Fig. 5A) to trigger RNAseH-dependent degradation of this lncRNA. We first evaluated the efficacy of two ASO candidates in vitro. We transfected the murine neuroblastoma cell line Neuro 2a (N2a) with either of the ASOs (Snhg11-ASO 1 and Snhg11-ASO 2) or with a negative control (Control-ASO) consisting in a gapmer of the same length as the ASOs with a scrambled sequence not targeting any gene. 72h after transfection, we observed a dramatic reduction of *Snhg11* expression as measured by RT-qPCR in cells transfected with ASO 2 (Fig. 5B) but not with ASO1. To establish the functional knockdown, as Snhg11 was previously reported to induce increased proliferation in different cancer cells[45-47], we analyzed whether its reduction impacted the proliferative capacity of N2a cells. The CCK8 assay revealed that the ASO2-mediated knockdown of Snhg11 dramatically reduces the proliferation of N2a cells, 24h after transfection and their proliferative capacity is arrested for at least 72h in a dose-independent manner (Fig. 5C and D). We did not observe any alteration of cell proliferation in cells transfected with ASO1 (Supplementary Fig. 7), in line with its lack of effectiveness in reducing Snhg11 expression. This lack of proliferation in the absence of Snhg11 was also reproduced in a Colony Formation Assay (CFA), showing that cells transfected with 50nM or 80nM but not 30nM ASO2 completely lose their capability to grow into a colony (Fig. 5E and F).

**Figure 5.**
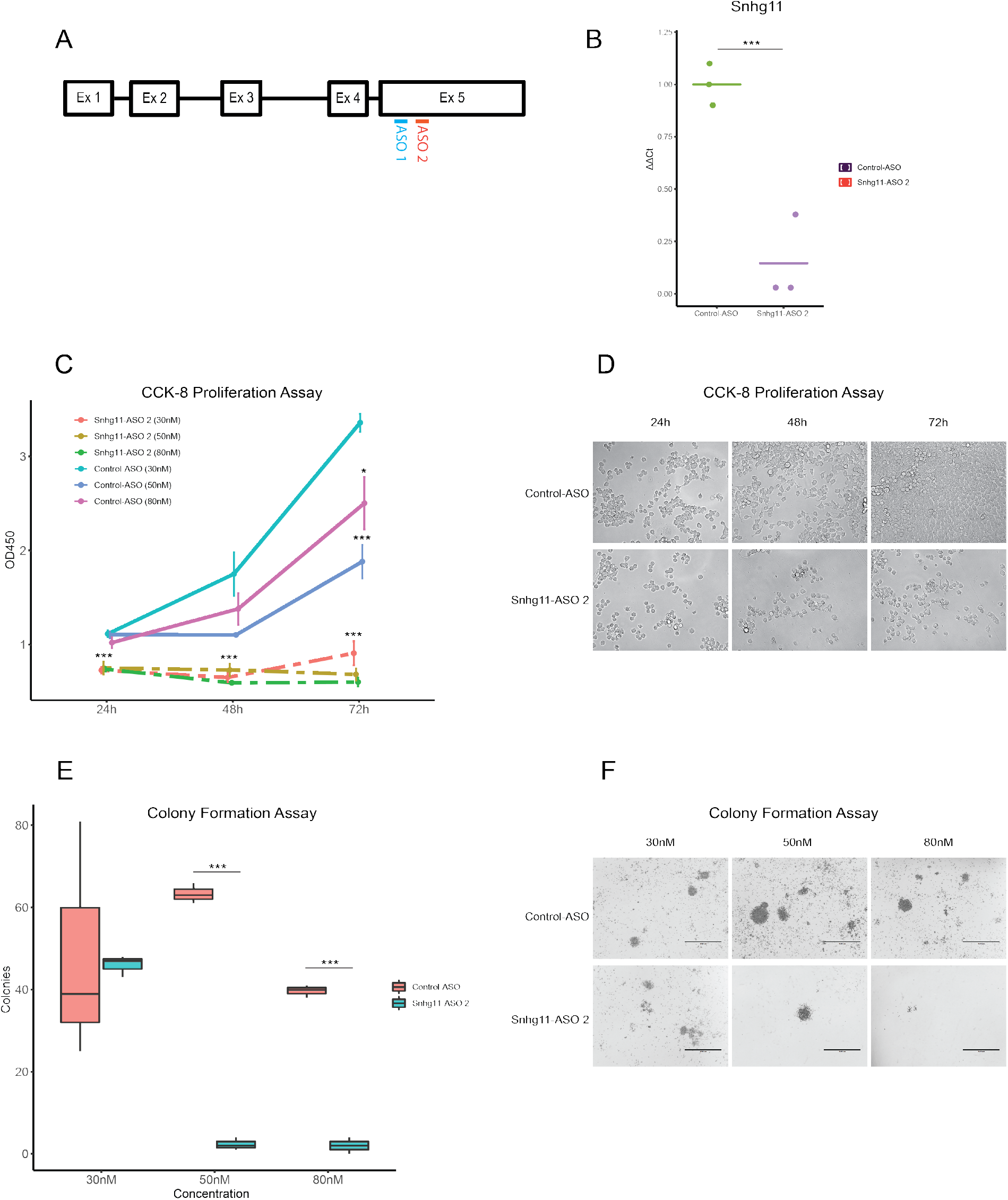
*In vitro* validation of ASO-mediated Snhg11 knockdown. **(A)** Exonic targets of the two ASOs designed. **(B)** RT-qPCR analysis of Snhg11 expression in N2a cells 72 hours after transfection. *** p<0.001; Student t-test. **(C)** CCK8 quantification of cell proliferation for 72 hours after transfection of NC or ASO. * p< 0.05, *** p<0.001; TukeyHSD test. **(D)** Representative images of the proliferative state of cells transfected either with 30nM control-ASO or with 30nM ASO 2 after 24h, 48h and 72h. **(E)** Colony count of transfected cells in the Colony Formation Assay. *** p<0.001; Student t-test. **(F)** Representative images of the N2a colonies 21 days after being transfected with increasing concentrations (30nM, 50nM and 80nM) of control-ASO or ASO 2.

### ASO-mediated knockdown of Snhg11 *in vivo* impairs memory and adult neurogenesis

Once the efficacy of the ASO-mediated knockdown of Snhg11 was confirmed in vitro, we investigated its impact in vivo by a bilateral injection of 50 ug of either ASO 2 or NC specifically in each of the hemispheres of the dorsal DG of 4 months-old WT mice (Fig. 6A). The specific knockdown of Snhg11 in the dorsal hippocampus was confirmed by RT-qPCR (Fig. 6B), to rule possible off-target effects (Supplementary Fig. 8A). Interestingly, the reduction in *Snhg11* expression was accompanied by a reduction in *Ntng1* expression, suggesting a possible crosstalk between the two genes (Supplementary Fig. 8B). In a second series of experiments, we performed a behavioural battery upon ASO or Control-ASO injection, to assess the relevance of Snhg11 for different types of memory (Fig. 6A). Strikingly, the reduction of Snhg11 had a dramatic impact on all the cortico-hippocampal memory paradigms tested. In the Novel Object Recognition (NOR), a task measuring recognition memory, non injected and WT mice injected with the Control-ASO properly acquired memory and recalled it both after 1 hour (Fig. 6C) or after 24 hours (Fig. 6D) as shown by the pronounced preference for the novel object measured by the Discrimination Index (DI). Instead, WT animals that were administered with ASO2 failed to display object recognition both at short or long term to a similar extent as trisomic mice (Fig. 6D). In order to address more specifically the impact of *Snhg11* knockdown, we assessed whether ASO-injected animals presented an altered spatial pattern separation memory, which strictly depends on the DG. For this, we used a paradigm known as object pattern separation [48]. In this case, the loss of Snhg11 led to an even more dramatic reduction of memory performance than the one observed in TS, whereas both WT and Control-ASO-injected mice showed a high DI (Fig. 6E), reflecting a well-functioning DG and hippocampal circuit. Interestingly, the impact of Snhg11 knockdown was specific to memory functions, as no changes were observed on motor activity, explorative behaviour or anxiety (Supplementary Fig. 7).

**Figure 6.**
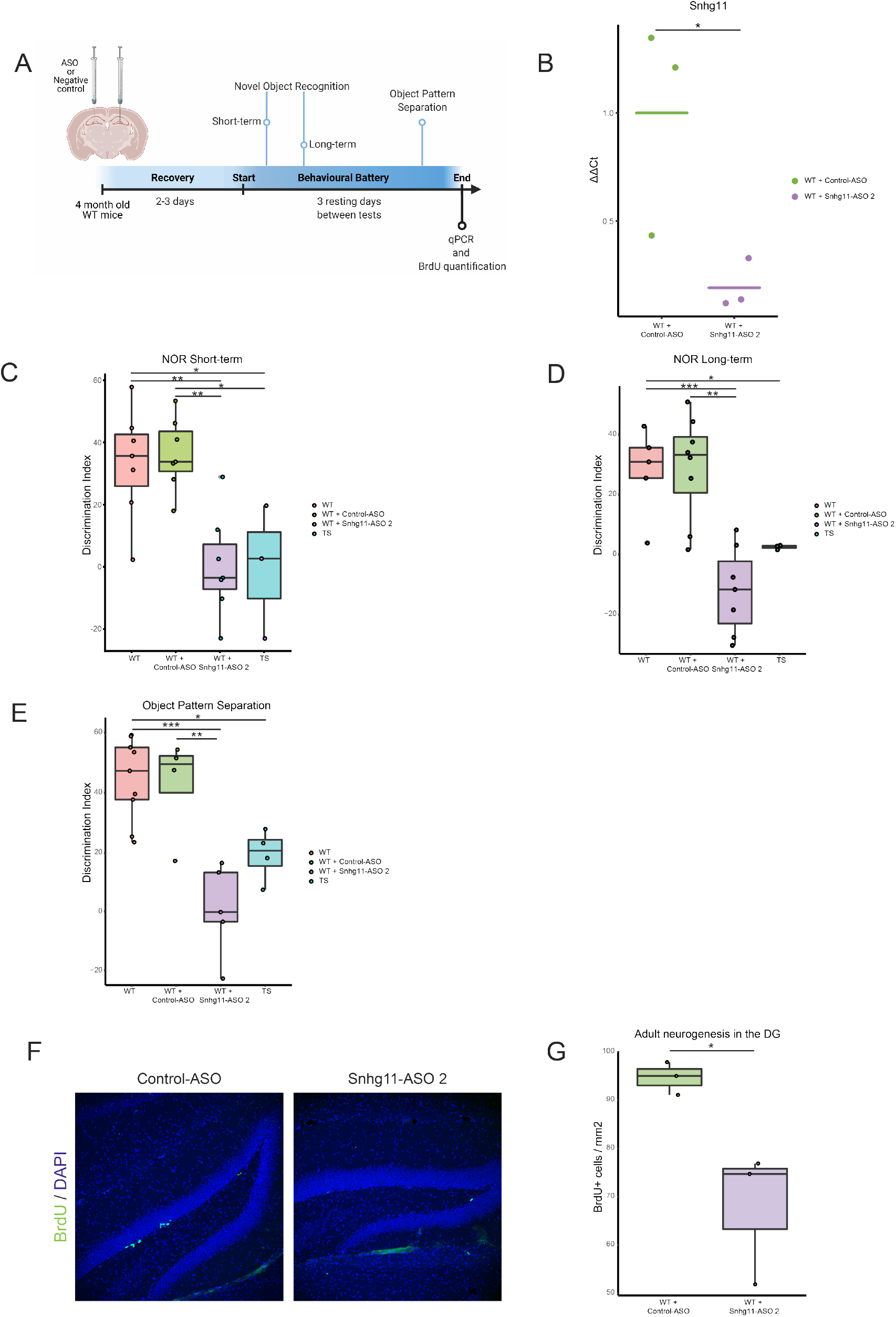
*In vivo* knockdown of *Snhg11* expression in the WT DG. **(A)** *Outline of the experimental plan*. **(B)** RT-qPCR analysis of Snhg11 expression in the dorsal hippocampus of injected animals. * p<0.05 Mann-Whitney-Wilcoxon test. **(C)** Novel Object recognition memory after 1 hour and after 24 hours **(D). (E)** Pattern separation memory after 1 hour. * p< 0.05, ** p<0.01,*** p<0.001; TukeyHSD test. **(F)** Representative images of BrdU labeled in sections of the dentate gyrus in a Control-ASO-injected mouse (left) and Snhg11-ASO 2-injected mouse (right). **(G)** Quantification of BrdU+ cells in the dentate gyrus. * p< 0.05; Mann-Whitney-Wilcoxon test.

## Discussion

Trisomy of HSA21 has been reported to produce a global perturbation of the transcriptome, showing not only an aberrant expression of genes located in the triplicated chromosome, but also a high number of differentially expressed genes (DEGs) mapping to all other chromosomes [19]. However, as the mechanisms regulating gene expression are highly cell-type specific, the full complexity of gene deregulation in DS may be missed in bulk studies. This is particularly relevant in those tissues with high cellular heterogeneity such as the hippocampus, a region particularly affected in DS, where the extent to which each cellular subtype is affected by the trisomy remains elusive.

To overcome this limitation, we generated the first single-nuclei atlas of the trisomic hippocampus by characterizing the transcriptome of tens of thousands of individual hippocampal neurons in parallel. Single-nuclei sequencing enabled us to characterize the nascent transcriptome of a large population of hippocampal neurons in euploid and trisomic mouse brains. From more than 50,000 nuclei, we built a high resolution gene expression atlas of the complex population of neurons that integrates the hippocampus both in WT and in a mouse model of DS, thereby unveiling cell-specific molecular alterations caused by the trisomy. In fact, we found almost half of the DEGs deregulated in a single neuronal type, thus indicating that the transcriptomic perturbations caused by the triplication are highly dependent on cellular identity. Our study also represents the first unbiased comprehensive assessment of the changes in the hippocampal neuronal composition that derive from the trisomy. Our data revealed abnormal cell composition specifically of the Ts65Dn dentate gyrus and of specific subtypes of interneurons, without major changes in CA1, CA2, CA3 and RHP.

A large body of evidence has shown that synaptic deficits and cognitive impairment in mouse models of DS largely derives from an altered excitatory-inhibitory balance [4, 36]. In fact, the number of GABAergic interneurons is increased in the hippocampus of TS [17, 18, 49]. However, many of these studies are limited to the quantification of specific neuropeptides that only represent a fraction of the total interneuron population in the hippocampus, and there have been contradictory results in regards to the specific subpopulations that are amplified. The assumption is that the increased interneuron population arises from an amplified neurogenesis during embryonic development of neural progenitor cells in the medial ganglionic eminence (MGE). Nevertheless, we found an increased population of neurogliaform cells derived from the CGE along with increased hippocamposeptal interneurons, which is a population with a high expression of somatostatin *Sst*+ neurons [38]. Instead, chandelier and basket cells, which derive from the MGE, and *Cck*-expressing interneurons, derived from the CGE, remained unaltered in the trisomic hippocampus. These results suggest that the amplification of the interneuron population in DS is not limited to an increased neurogenesis in the MGE but also in the CGE. We also quantified the number of nuclei in inhibitory clusters expressing classical interneuron neuropeptides. We found Sst to be the most clearly overrepresented neuropeptide in the trisomic hippocampus, which is consistent with previous reports in CA1 [18]. Similarly to Hernandez *et al*. [17], we also found an increased population of *Vip*+ neurons, while the *Pvalb*+ population remained unchanged. However, no changes were observed in *Npy, Calb* and *Calb2* populations.

The single-nuclei approach allowed us to identify Kcna1 and Kcnc1, encoding two voltage-gated K+ channel subunits, Kv1.1 and Kv3.1, to be downregulated specifically in the inhibitory population. The Kv1 family of voltage-gated K+ channel raises the action potential (AP) firing threshold in a given neuron [50], while the Kv3 family accelerates AP repolarization [51]. Thus, downregulation of these subunits would lead to an increased excitability specifically in trisomic interneurons, thereby possibly contributing to the excitatory/inhibitory imbalance that contributes to cognitive deficits in TS. Our results support the hypothesis that the increase of spontaneous GABAergic events observed in TS mice [18] may be due to an enhancement in interneuron excitability rather than an increase in the number of specific GABAergic interneurons and synapses [36].

The second interesting finding was the reduced population of granule cells in dentate gyrus (DG) of Ts65Dn mice. In this region, a relatively small alteration in the number of granule cells might have an impact in learning and memory [52]. The granule cell hypocellularity observed in Ts65Dn mice has been suggested to be contributed by altered neurogenesis in DS, and in our study we detected that the DEG of the trisomic DG are specifically enriched in genes related to neurogenesis such as *Flrt3* and *Ptprd*, which could contribute to explain the defective adult neurogenesis observed in DS mouse models [14, 40, 41]. Strikingly, the DG showed a specific shift in the UMAP representation of the nuclei transcriptome. Although this shift did not translate into an enrichment of DEGs, this region showed a significant loss of expression of genes enriched in the DG according to the Allen Brain Atlas [33]. In particular, *Ntng1*, a marker of mature granule cells [53], was consistently downregulated in the trisomic DG. We also detected a marked reduction of the expression of *Snhg11*, a lncRNA described to be involved in cell proliferation in different types of cancer [45-47], mainly through the regulation of the Wnt/β-catenin signaling pathway, which is affected in DS [54]. Despite its high expression in the murine hippocampus, the role of this lncRNA in neuronal functions has not been explored. Here, we investigated the impact of the reduction of *Snhg11* using an ASO in wild type mice to resemble the situation detected in the Ts65Dn hippocampus. Upon *in vitro* knockdown of *Snhg11* in neuroblastoma cells, we observed a dramatic reduction of cell proliferation, that translated to a reduced adult neurogenesis after ASO-mediated *Snhg11* knockdown in the adult WT DG. As expected, due to the critical role of adult neurogenesis in long-term memory [55, 56], we detected a strong impairment of long-term memory in the NOR task upon upon *Snhg11* reduction. However, we also observed a defective short-term memory in the NOR task and in the DG-specific object pattern separation task. Although adult neurogenesis importance for short-term memory remains controversial [55, 57], it has to be borne in mind that small nucleolar host genes (SNHGs) have wide-range mechanisms of action [44] that could influence the function of granule cells, thereby contributing to DG-dependent memory deficits. In fact, we showed that the expression on this lncRNA is not limited to the nucleus, but is also expressed in the cytoplasm, where it could act as a sponge for miRNAs, bind mRNAs to repress translation or prevent protein ubiquitination, as many members of the same family [44]. Interestingly, the knockdown of *Snhg11* by itself was sufficient to induce the downregulation of *Ntng1*.

Here, we performed the first comprehensive assessment of the cellular and molecular changes that occur in the hippocampus of a mouse model of DS, thereby identifying genes, processes and cell types impacted by the trisomy. These results provide evidence of the high cell-type specificity of the transcriptomic alterations previously described in DS. Furthermore, we identified *Snhg11* as a key player in DG function, with a role on cell proliferation, adult neurogenesis and memory formation, and show that its downregulation mimmicks DS phneotypes, and could thus contribute to the DG-dysfunction observed in DS.

## Supporting information

Supplementary table 1: cluster markers

Supplementary table 2: interneuron subtypes markers

Supplementary table 3: DEGs

Supplementary table 4: ASO sequences

## Author contributions

MD and CS conceived the study, designed and coordinated the study, and wrote the manuscript. CS collected and analyzed sequencing datasets. CS conducted animal and in vitro experiments. IdT applied bootstrap analyses.

## Materials and Methods

### Animals

Experimental mice were generated by crossing of Ts65Dn females to C57/6Ei × C3H/HeSnJ F1 hybrid (B6EiC3) males. The parental generation was obtained from the research colony at the Jackson Laboratory (B6EiC3Sn a/A-Ts(1716)65Dn/J; Stock No: 001924). Genotypes of mice were authenticated by PCR assays on mouse tail samples with an in-house protocol. Mice were housed in standard cages (156 × 369 × 132 mm), with food and water available ad libitum in standard conditions (12:12 light cycle; 400 lux). Sawdust and nesting materials in each cage were changed once a week, but never on the day before or the day of testing, to minimize the disruptive and stressful effect of cage cleaning on behavior.

All experiments followed the principle of the “Three Rs”: replacement, reduction and refinement according to Directive 63 / 2010 and its implementation in Member States. The study was conducted according to the guidelines of the local (law 32/2007) and European regulations (2010/63/EU) and the Standards for Use of Laboratory Animals no. A5388-01 (NIH), and approved by the Ethics Committee of Parc de Recerca Biomèdica (Comité Ético de Experimentación Animal del PRBB (CEEA-PRBB); MDS 0035P2). The CRG is authorized to work with genetically modified organisms (A/ES/05/I-13 and A/ES/05/14).

### Single Nucleus RNA sequencing

#### Nucleus isolation

Mice were sacrificed and the hippocampus of each mouse was dissected and placed in chilled Hanks’ Balanced Salt Solution (Sigma #55021C). The “Frankenstein” protocol [ref] was followed to obtain a nuclei suspension. Each hippocampus was transferred to a new tube containing 500μL chilled EZ Lysis Buffer (Sigma #3408) and homogenized using a sterile RNase-free douncer (Mettler toledo #K-749521-1590). The resulting homogenate was filtered using a 70μm-strainer mesh to remove remaining chunks of tissue and centrifuged at 500g for 5 minutes at 4ºC. The pellet containing the nuclei was resuspended in 1.5mL EZ Lysis Buffer and centrifuged again. The supernatant was removed and 500μL of Nuclei Wash and Resuspension Buffer (NWRB, 1X PBS, 1% BSA and 0.2U/μL RNase inhibitor (Thermo Scientific #N8080119) were added without disturbing the pellet and incubated for 5 minutes. After incubation, 1mL of NWRB was added and the pellet resuspended. The nuclei suspension was centrifuged again and the washing step was repeated with 1.5mL of NWRB. After an additional centrifugation, nuclei were resuspended in 500μL of 1:1000 anti-NeuN antibody conjugated with AlexaFluor 647 (ab190565 Abcam) in PBS and incubated in rotation for 15 minutes at 4ºC. After incubation, nuclei were washed with 500μL of NWRB and centrifuged again. Last, nuclei were resuspended in NWRB supplemented with DAPI and filtered with a 35μm cell strainer to obtain a single-nuclei suspension.

#### 10x single-cell barcoding, library preparation and sequencing

Fluorescent activated nuclear sorting (FANS) was used to sort NeuN+ neuronal nuclei. Using a 70um nozzle to minimize the volume deposited, 10.000 nuclei from each sample are sorted directly into a 96-well plate prefilled with 10X RT buffer prepared without the RT Enzyme Mix. After sorting, the RT Enzyme C is added and the volume of each well topped up to 80ul with nuclease free water. Last, 75uL of the nuclei plus RT mix are loaded into the Chromium Single Cell Chip. All downstream cDNA synthesis), library preparation, and sequencing were carried out as instructed by the manufacturer (10x Genomics Chromium Single Cell Kit Version 3). Libraries were prepared and sequenced in two separate experiments. In the first three wild type and three trisomic hippocampi and in the second one hippocampus per genotype were processed. For each experiment, all libraries were pooled and sequenced on NovaSeq 6000 S1 to an average depth of approximately 20.000 reads per cell.

#### 10X data pre-processing

The resulting reads were aligned to the reference genome and converted to mRNA molecule counts using the Cellranger pipeline (CellRanger v3.0.1) provided by the manufacturer. For every nucleus, we quantified the number of genes for which at least one read was mapped, and then excluded all nuclei with fewer than 200 or more than 2500 detected genes, to discard low quality nuclei and duplets, respectively. Genes that were detected in fewer than six nuclei were excluded. Expression values Ei,j for gene i in cell j were calculated by dividing UMI counts for gene i by the sum of the UMI counts in nucleus j, to normalize for differences in coverage, and then multiplying by 10,000 to create TP10K (transcript per 10,000) values, and finally computing log2(TP10K + 1) (using the NormalizeData function from the Seurat package v.2.3.4 [58].

#### Batch Correction and scaling data matrix

Since samples were processed in two different experiments, batch correction was done using Harmony (ref) on the normalized dataset. The batch-corrected data was scaled using the ScaleData function from Seurat [ref] with default parameters (v.2.3.4), yielding the relative expression of each gene by scaling and centering. The scaled data matrix was then used for dimensionality reduction and clustering. To rule out the possibility that the resulting clusters were driven by batch or other technical effects, we examined the distribution of samples within each cluster and the distribution of the number of genes detected across clusters (as a measure of nucleus quality). Overall, the nuclei separated into distinct point clouds in t-distributed stochastic neighbor embedding (t-SNE) space that were not driven by batch; each cluster was a mixture of nuclei from all technical and biological replicates.

#### Dimensionality reduction, clustering and visualization

We used the scaled expression matrix restricted to the variable genes for principal component analysis (PCA), using the RunPCA method in Seurat (a wrapper for the irlba function), computing the top 60 principal components. Scores from these principal components were used as the input to downstream clustering and visualization by Uniform Manifold Approximation and Projection (UMAP).

Clustering was performed using the Seurat functions FindNeighbors and FindClusters (resolution = 0.6). Clusters were then visualized with UMAP. Reference anchors were identified between genotypes before integration with the IntegrateData function, and integrated data were then processed by the same methods.

#### Identification of marker genes of individual cell clusters

Cluster-specific marker genes were identified using the FindAllMarkers function, utilizing a negative binomial distribution (DESeq2). A marker gene was defined as being >0.25 log-fold higher than the mean expression value in the other clusters, and with a detectable expression in > 20% of all cells from the corresponding cluster. In this way, we were able to select markers that were highly expressed within each cluster, while still being restricted to genes unique to each individual cluster.

#### Identification of DEGs between WT and TS

All clusters identified as belonging to the same cell type were merged for differential gene expression analyses. Within each cell type, WT and TS samples were compared for differential gene expression using Seurat’s FindMarkers function. To be included in the analysis, the gene had to be expressed in at least 10% of the cells from one of the two groups for that cell type and there had to be at least a 0.1 fold change in gene expression between genotypes. After correcting for multiple testing, only genes with a p adjusted value < 0.001 were considered for downstream analyses.

#### Gene set enrichment

The differential expression signatures from each cellular subtype were tested for enriched Gene Ontology processes, using a hypergeometric test (function enrichGO from de clusterProfiler package in R[59]), and corrected for multiple hypotheses by FDR. Processes with p adjusted value < 0.05 were reported as significantly enriched. The complete list of genes present in the dataset was used as the universe for the hypergeometric test.

#### Diffusion map

The diffusion components were calculated using the cell embedding values in the top 15 principal components (generated either on the scaled expression matrix restricted to the variable genes in the 7-month-old mouse dataset or on the aligned canonical correlation analysis (CCA) subspace for the entire time-course data), using the DiffusionMap function from the destiny package [60] in R (with k = 30 and a local sigma). We then chose the top two diffusion components for data visualization. To investigate the DG-specific shift, we took advantage of the gene_overlap [61] function of the destiny package, that allows us to identify drivers of cell embedding at a global or subregion level from non-linear low-dimensional embeddings such as UMAP or Diffusion Map.

#### Cellular proportion analysis

To gain insight into the cell type alterations in the trisomic hippocampus, the relative proportion of the number of nuclei in each cell type was normalized to the total number of nuclei captured from each library. To determine if any changes in cell-type proportion were statistically significant, we implemented single cell differential composition analysis (scDC) [34]to bootstrap proportion estimates for our samples. We employed a linear mixed model (random effect of subject) to determine if any changes in cell-type proportion were present.

#### RNAscope

In situ hybridization was performed using RNAscope® Multiplex Fluorescent assay (V1) (Advanced Cell Diagnostics) probes and reagents. Naive animals were sacrificed and transcardially perfused with 50mL of chilled PBS followed by fixation with 50-100mL of 4% depolymerised paraformaldehyde (PFA). Following hippocampus dissection, tissues were placed in 4% PFA and post-fixed overnight at 4ºC. Tissues were then immersed in increasing concentrations of sucrose (10%, 20% and 30%) over three days, increasing the concentration every 24 hours. Once embedded in sucrose, tissues were dried and frozen at -80ºC prior to cryosectioning. 14 μm sections were obtained and mounted on Superfrost charged slides and stored at - 80ºC until being processed. In situ hybridization was performed following the manufacturer’s guidelines. Custom RNAscope® target-specific oligonucleotide (ZZ) probes were designed by Advanced Cell Diagnostics targeting nucleotides 2-1604 for Snhg11 (NR_164123.1) and 1130 - 2109 for Ntng1 (NM_030699.2).

### ASO-mediated knockdown of Snhg11

#### Antisense oligonucleotides

ASOs (IDT) were uniformly modified with 2′-O-(2-methoxy) ethyl sugars (2′MOE), phosphorothioate backbone, and 5′-methyl cytosine. ASOs were dissolved in 0.9% saline to a concentration of 75μg/μL. ASO sequences are provided in Supplementary Table 4. Both ASO sequences were subjected to a BLAST search. The Snhg11 ASOs 1 and 2 had only positive matches for the target sequences and no other mouse coding sequence. The control ASO (Control-ASO), which included the same nucleotides in a scrambled order, did not generate any full match to identified gene sequences in the database.

### In vitro knockdown of Snhg11

#### Cell culture and ASO transfection

Mouse N2a neuroblastoma cells were maintained in high glucose Dulbecco’s modified Eagle’s medium (Thermo Fisher #11965084) supplemented with 10% FBS, 100U/mL penicillin and 100ug/mL streptomycin in a humid atmosphere containing 95% air and 5% CO2 at 37ºC. ASOs and NC were transfected into N2a cells using Lipofectamine 3000 (Thermo Fisher #L3000001) in a 96-well format. For each well, 1.5μL of the ASO or NC were diluted in 8.5uL Opti-MEM medium (Thermo Fisher #31985062), 0.3μL P3000 and 0.25μL Lipofectamine and incubated at room temperature for 15 minutes. Meanwhile, N2A cells were trypsinized, counted and diluted to 22000 cells/140 μL with antibiotic-free DMEM supplemented with 10% FBS. After incubation, 10μL of the ASO-lipid complex was added to each well in triplicate followed by 140μL of the diluted cells for a final volume of 150 μL.

#### RNA extraction from N2a

To ensure Snhg11 knockdown, 72 hours after being transfected, N2a cells were washed with pre-warmed PBS and trypsinized. Cells were then pelleted at 300g for 7 minutes and resuspended in 200μL of Homogenization Buffer from the Maxwell RSC simplyRNA Tissue Kit (Promega #AS1340). Cells were then lysed by adding 200μL of lysis buffer and transferred into the Maxwell RSC Cartridge. Cartridges were then loaded into the Maxprep Liquid Handler and RNA was extracted in 30 μL of water following the simplyRNA Cells protocol provided by the manufacturer.

#### Quantitative Real-time PCR

All experiments were performed with three biological and three technical replicates. Briefly, RT-qPCR reactions were carried out using LightCycler® 480 SYBR Green I Master Mix (Roche #04-887-352-001) on a LightCycler® Real-Time PCR System (Roche). The final volume for each reaction was 20 μl with 500 nM of corresponding gene specific primers (Snhg11 forward primer: ctgtccagctaggaagcagc, Snhg11 reverse primer: tcgctccacactgatgttgg, Ntng1 forward primer: aggggcaagagaccaagg, Ntng1 reverse primer: agggatggtgtctatcgtcct, Gapdh forward primer: ggagattgttgccatcaacga, Gapdh reverse primer: tgaagacaccagtagactccacgac), and 5 μl of total cDNA diluted 1:5. A negative water control was included in each run. The thermal cycling was initiated at 95 °C for 10 minutes followed by 45 cycles of 10 s at 95 °C and 15 s at 55ºC, the optimal annealing temperature for our target genes. Melting curve analyses were carried out at the end of each run of qPCR to assess the production of single, specific products.

#### CCK8 proliferation assay

ASO and NC transfected N2a cells were seeded into 96-well plates overnight. At 24, 48, or 72 h post-transfection, 10 μl Cell Counting Kit-8 solution (MedChem Express HY-K0301) was added into each well of 96-well plates, followed by further incubation of 3 h at 37 °C. Then the absorbance was determined at a wavelength of 450 nm to assess cell proliferation.

#### Colony Formation Assay

ASO and NC transfected cells were plated in 6-well plates at 500 cells/well in triplicate. The cells were maintained in cell culture with media change every 3 days. After 21 days, they were washed 3 times with PBS, fixed with 4% PFA for 10 minutes, stained with a 0.06% crystal violet solution for 1 hour at 37ºC and colonies counted under an inverted microscope.

### *In vivo* knockdown of Snhg11

#### Stereotactic infusion

For in vivo injection of ASOs, 4 month old male (n =7 per condition) were anesthetized by intraperitoneal injection of ketamine (75mg/kg) and meta (10mg/kg) and placed on a stereotaxic frame. For each injection, a small incision was made, the skull was exposed, and a small burr hole was drilled at the proper coordinates (antero-posterior, -2.2 mm ; mediolateral, +/-1.3 mm; and dorsoventral, −2 mm relative to the bregma). Three minutes after the needle (Hamilton #65458-01) was placed into the proper coordinates, a total of 660nL of saline-diluted ASO or NC at a concentration 75μg/μl was delivered into each hemisphere of the dentate gyrus at an infusion rate of 50nL/minute using an injection pump. After the infusion was completed, the needle was kept in the same position to ensure no contamination of other brain areas with residual volume. The incision site was sutured using surgical glue (Cemave #1050052) and the mouse was allowed to recover in a temperature-controlled environment. Following surgery, buprenorphine was administered at 0.1mg/kg/day and for 3 days. During this time, weight, grooming activity and home cage activity were controlled. In the first round of experiments, the site of injection in the dorsal DG was confirmed by the injection of 1μL of trypan blue in anesthetized mice.

#### Hippocampi isolation, RNA extraction and cDNA synthesis

To ensure knockdown of Snhg11, the hippocampus was dissected, snap-frozen in liquid nitrogen and kept at -80ºC until processed. Prior to RNA extraction, hippocampi samples were homogenized by mechanical trituration with a pestle. Total RNA extraction was performed using the RNeasy Mini Kit (Qiagen #74104) according to the manufacturer’s instructions. RNA purity was assessed using a NanoDrop 2000 (Thermo Scientific #ND-2000). RNA concentrations ranged from 30 to 200 ng/μl. All RNA samples were subjected to DNase digestion with DNase I (Sigma #AMPD1-1KT). Single-stranded cDNA was synthesized from 1000 ng of total RNA using SuperScript™ III Reverse Transcriptase and oligo(dT) primers (Thermo Scientific #18080093) following the manufacturer’s instructions. Quantitative Real-time PCR was performed as described above.

### Hippocampal dependent memory tests

#### Novel Object Recognition

The novel object recognition test (NOR), is a relatively fast and efficient means for testing different phases of learning and memory in mice. The test was conducted as explained in previously [62]. Briefly, the NOR protocol consists of four phases, namely habituation, familiarization, short-term test and long-term sessions. During the habituation session, mice were allowed to freely explore the empty open field for 10 minutes. 24 hours later, each mouse was returned to the arena containing two identical objects placed at symmetrical positions 5 cm from the arena wall and allowed to explore them freely for 15 minutes. During the familiarization session most mice reached a minimum exploration for each object of 30 seconds. Mice not reaching this criterion at 15 min, were excluded from the analysis, as it cannot be confirmed they spent enough time exploring to learn/discriminate. After a retention interval of 1 hour, the mouse was returned to the arena in which one of the objects was replaced by a novel object and let to explore both the familiar and the novel object for a total of 5 minutes. Similarly, 24 hours later the animal was returned to the arena in which the novel object of the short-term session was replaced by a third object. The animal was then let to explore both objects for 5 minutes. Time spent exploring each object in the three sessions was recorded.

#### Object Pattern Separation

To test pattern separation memory, the protocol described by van Hagen et al. (2015) [48], was followed with few modifications. The same square chamber used for NOR was used for OPS. Two days after completing the NOR, mice were habituated again to the open field without any objects for 10 minutes. 24 hours later, mice were presented with two identical objects placed in two symmetrical spots 5 cm from the arena wall. The objects were made with building blocks stuck to the arena floor so that mice were not able to move them. Animals were left 6 minutes to explore. The test session was performed 1 hour later, where one of the objects was placed in a novel location inside the arena 8 cm away from the initial location. Mice were left 4 minutes to explore. Time spent exploring each object in both sessions was recorded.

#### Behavioural experimental variables and data analysis

Locomotor activity was quantified as the total distance travelled in the apparatus during the experimental sessions. Thigmotaxis refers to the disposition to remain close to the walls of the apparatus. It is measured as distance travelled or percentage of time spent in the periphery of the apparatus. It decreases gradually during the first minutes of exploration, and can be used as an index of anxiety [63][ref]. Exploration time is defined as the action of pointing the nose toward an object, at a maximum distance of 2 cm or touching it. Going around the objects or sitting on the object is not evaluated as exploration time. As such, the exploration time is only computed when the snout of the animal is directed toward the object, sniffing or touching it. Discrimination index (DI), is the most relevant parameter in the NOR and OPS, which is a measure of the recognition memory of the animal. It is calculated as the difference in exploration time for the familiar versus the novel object in the case of NOR, or the familiar position versus the novel position in the OPS, divided by the total amount of exploration.

#### BrdU assay

Adult neurogenesis was measured by the incorporation of the thymidine analogue BrdU, which is incorporated into the DNA of dividing cells in detectable quantities during the S phase of cell division. After completing the behavioural battery, each animal received four intraperitoneal injections of BrdU (Merck #B5002) at 50mg/kg every 2 hours on a single day. Animals were sacrificed 24 hours after the last injection and transcardially perfused with 50mL of chilled PBS followed by fixation with 50-100mL of 4% (PFA). After perfusion, brains were removed and post-fixed overnight in 4% paraformaldehyde at 4°C. Sections 40 μm thick were then prepared in the coronal plane using a vibratome. From each animal, every sixth section (240 μm apart) of the series was used for BrdU Immunofluorescence. Sections were rinsed three times with PBS and incubated in 2M HCl for 15 minutes at 37ºC to denature DNA and allow the access for the anti-BrdU antibody. The acid was then neutralized by rinsing the sections with PBS. Sections were incubated with blocking solution (10% Donkey serum in TBS-Tween 0.5%) for 1h at room temperature and with the BrdU primary antibody (1:300; MBL, MI-11-3) diluted in blocking solution for 24h at 4ºC on a shaker. Sections were rinsed again with PBS and incubated with the secondary antibody (1:500; Donkey anti-mouse 488, Jackson #715-545-150) diluted in an incubation buffer (TBS-Tween 0.5% with 5% Donkey serum) for 3 hours at room temperature on a shaker. Rinsed sections were then mounted on uncoated Superfrost slides and covered with antifade and coverslip. For visualization and photography, specimens were observed under a confocal SP5 microscope.

## Acknowledgements

We would like to thank the Single Cell Genomics Team at the National Center for Genomic Analysis (CNAG) led by Holger Heyn for their assistance with experiment planning and sequencing and for the fruitful discussion throughout this project. This paper was typeset with the bioRxiv word template created by @Chrelli: www.github.com/chrelli/bioRxiv-word-template. Figure 1A was created with BioRender.com.

## Supplementary Figures

**Supplementary Figure 1.**
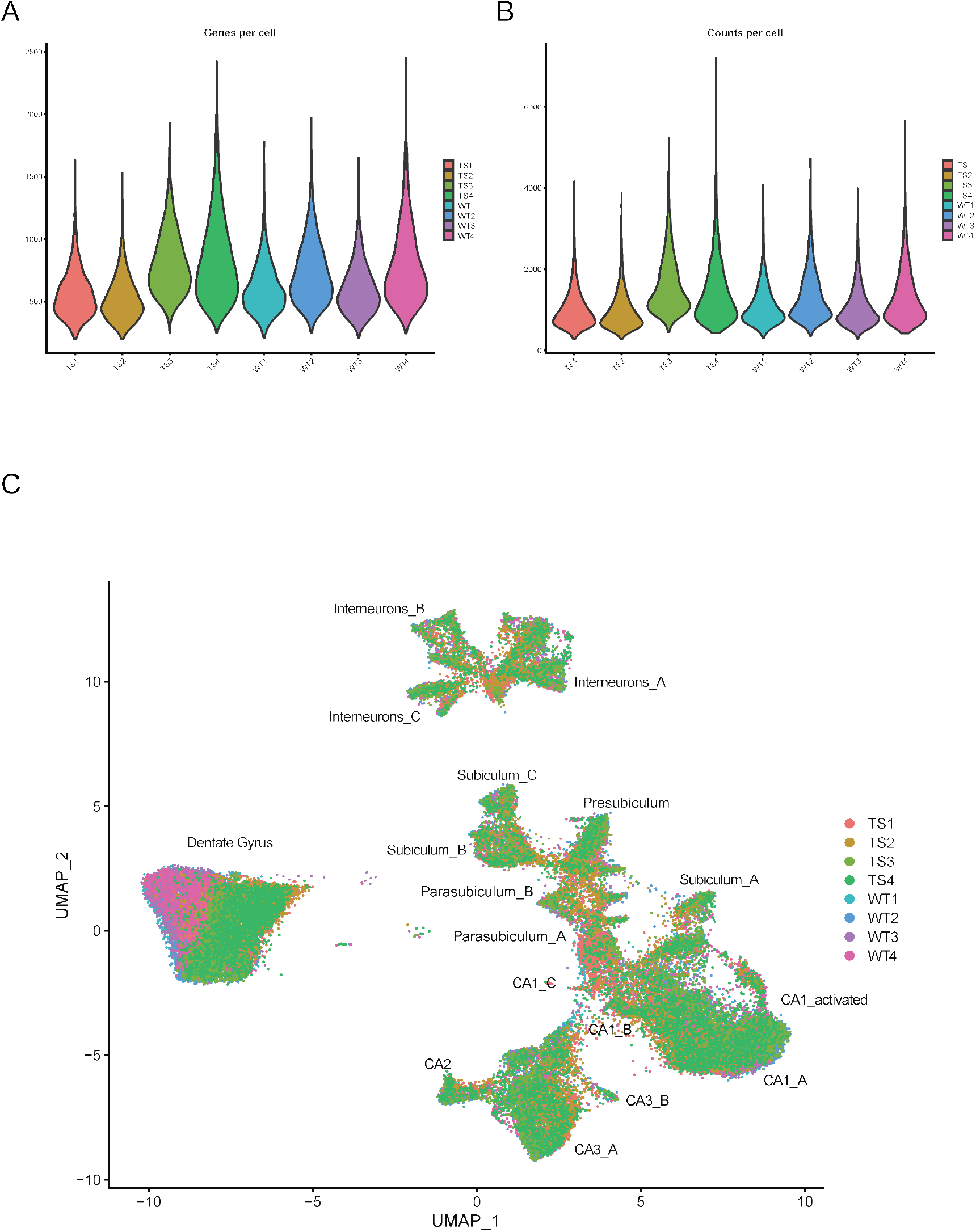
10x library quality control features. **(A)** Violin plot for the number of unique genes detected per cell in each. **(B)** *Violin plot for the number of unique transcripts detected per cell in each sample*. **(C)** Sample batch effects on cell clusters. Cells are colored by sample of origin on the UMAP plot. All the clusters are composed of a mixture of cells that originated from each of the eight samples.

**Supplementary Figure 2.**
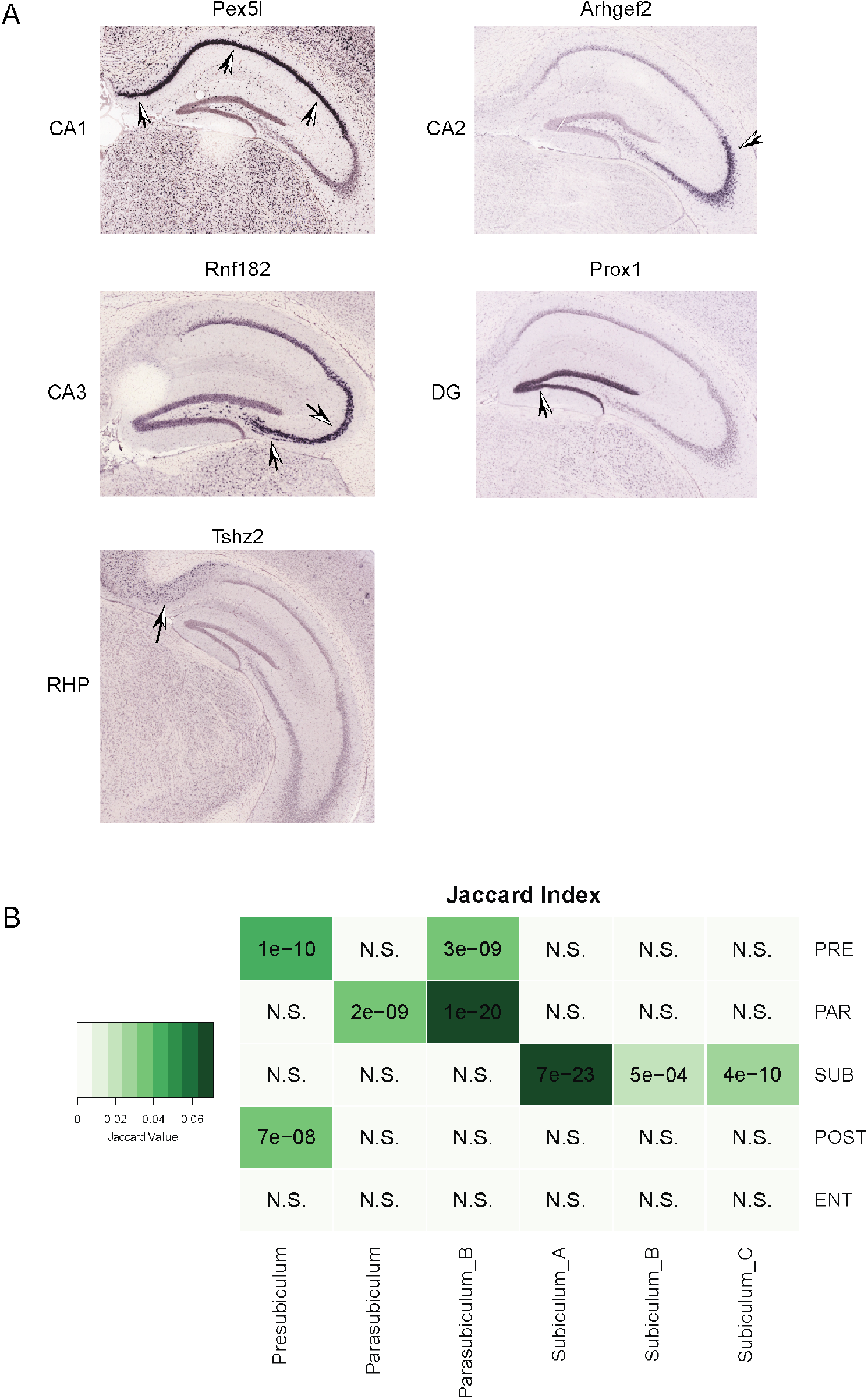
Validation of clusters identity. **(A)** Cross-validation of marker genes for specific hippocampal subpopulations identified in our dataset with the ISH images from the Allen Brain Atlas. **(B)** Jaccard overlap between cluster markers and enriched genes in each retrohippocampal subregion according to Allen Brain Atlas. p-values are indicated

**Supplementary Figure 3.**
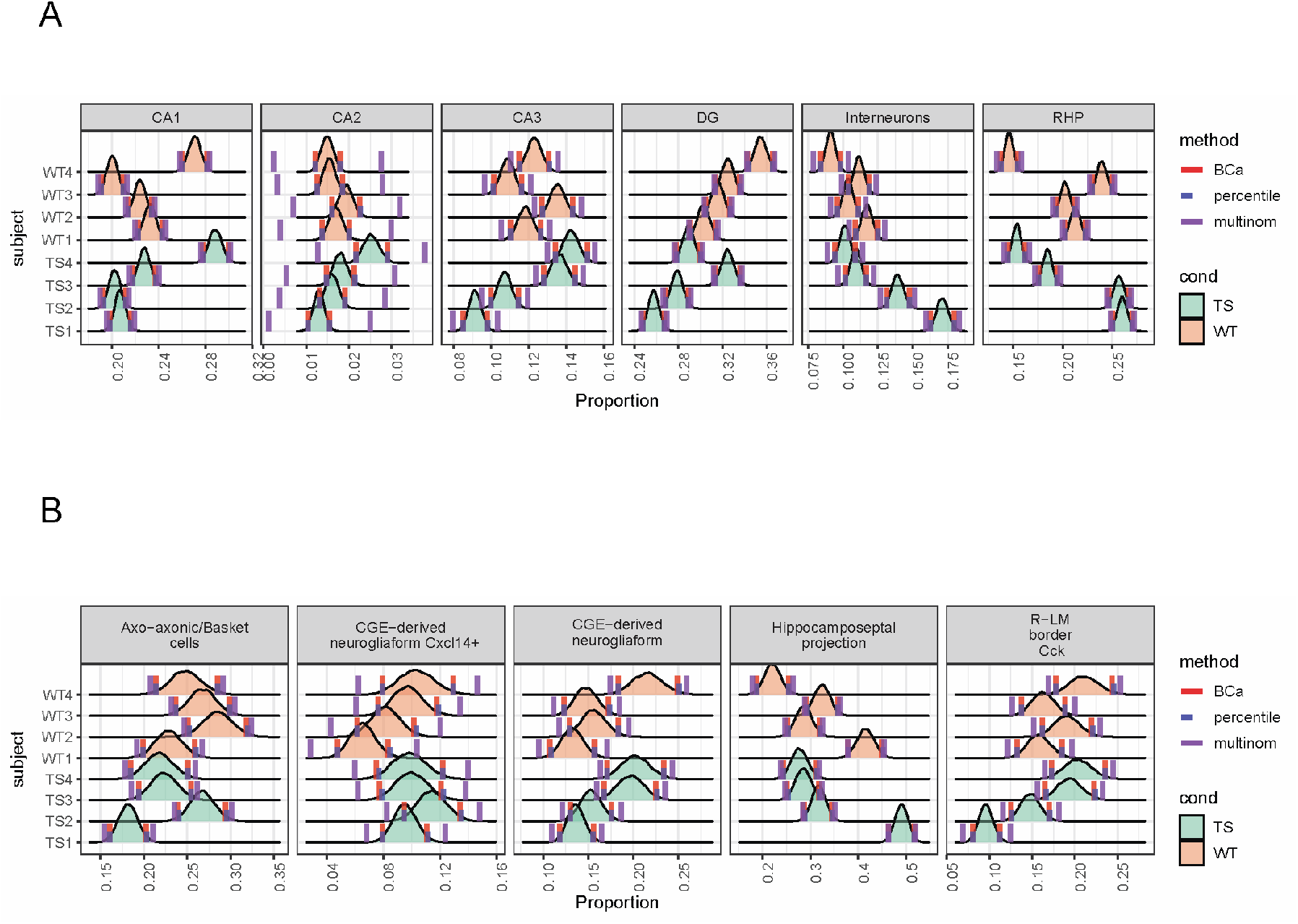
Single cell differential composition analysis. **(A)** Bootstrap estimated distribution of major hippocampal neuronal types and CA subregion subtypes for each sample. RHP: retrohippocampal neurons. **(B)** Bootstrap estimated distribution of the five identified interneuron subtypes. Confidence intervals were estimated with three different methods: multinomial proportions (purple line), bootstrap percentile method (blue line) and bootstrap BCa method (red line).

**Supplementary Figure 4.**
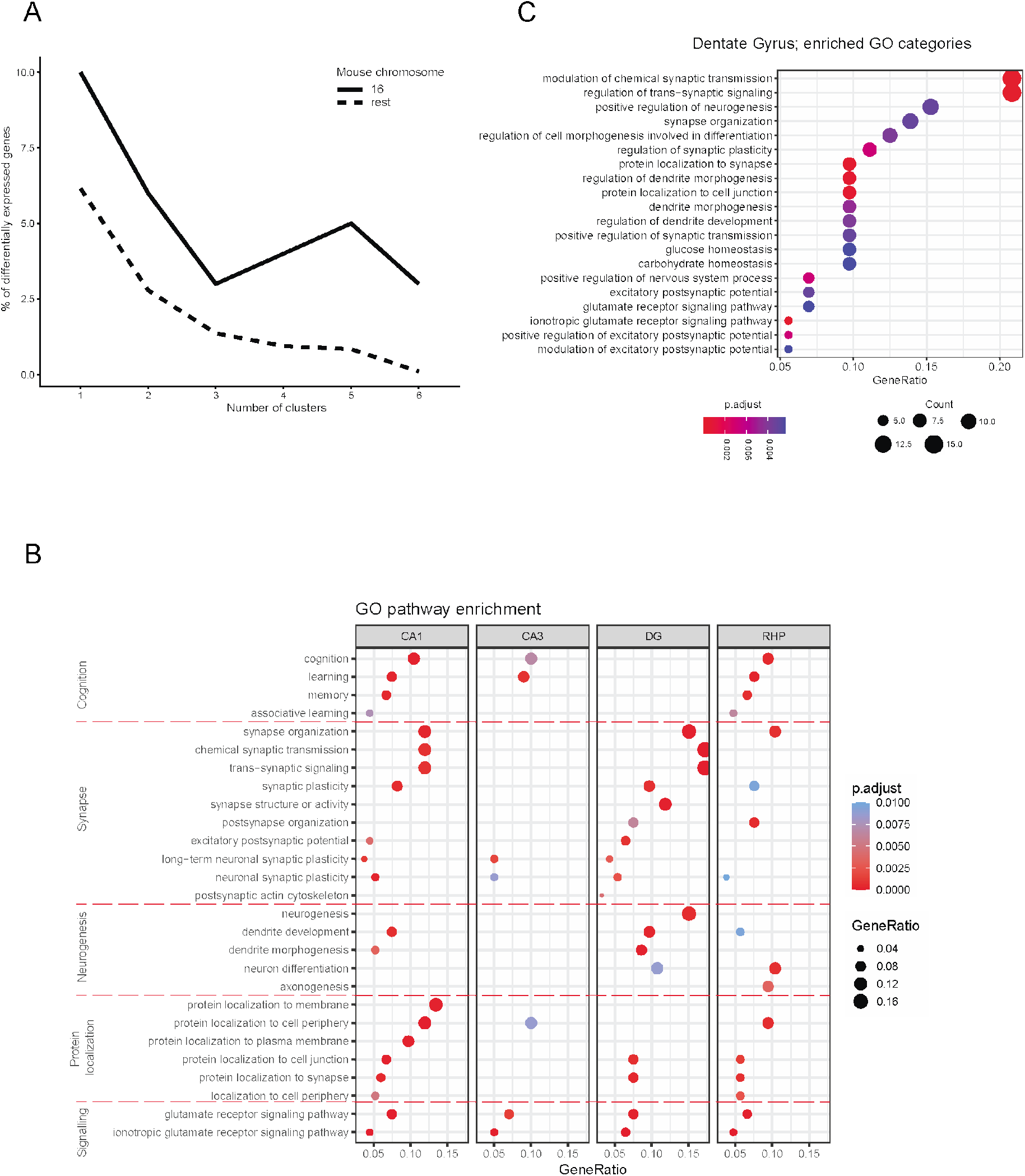
**(A)** Proportion of chromosome 16 or non-chromosome 16 DEGs altered in an increasing number of clusters. **(B)** Selected enriched gene ontologies (GOs) from DEGs in each major hippocampal subregion. **(C)** Enriched GO categories for DG DEGs

**Supplementary Figure 5.**
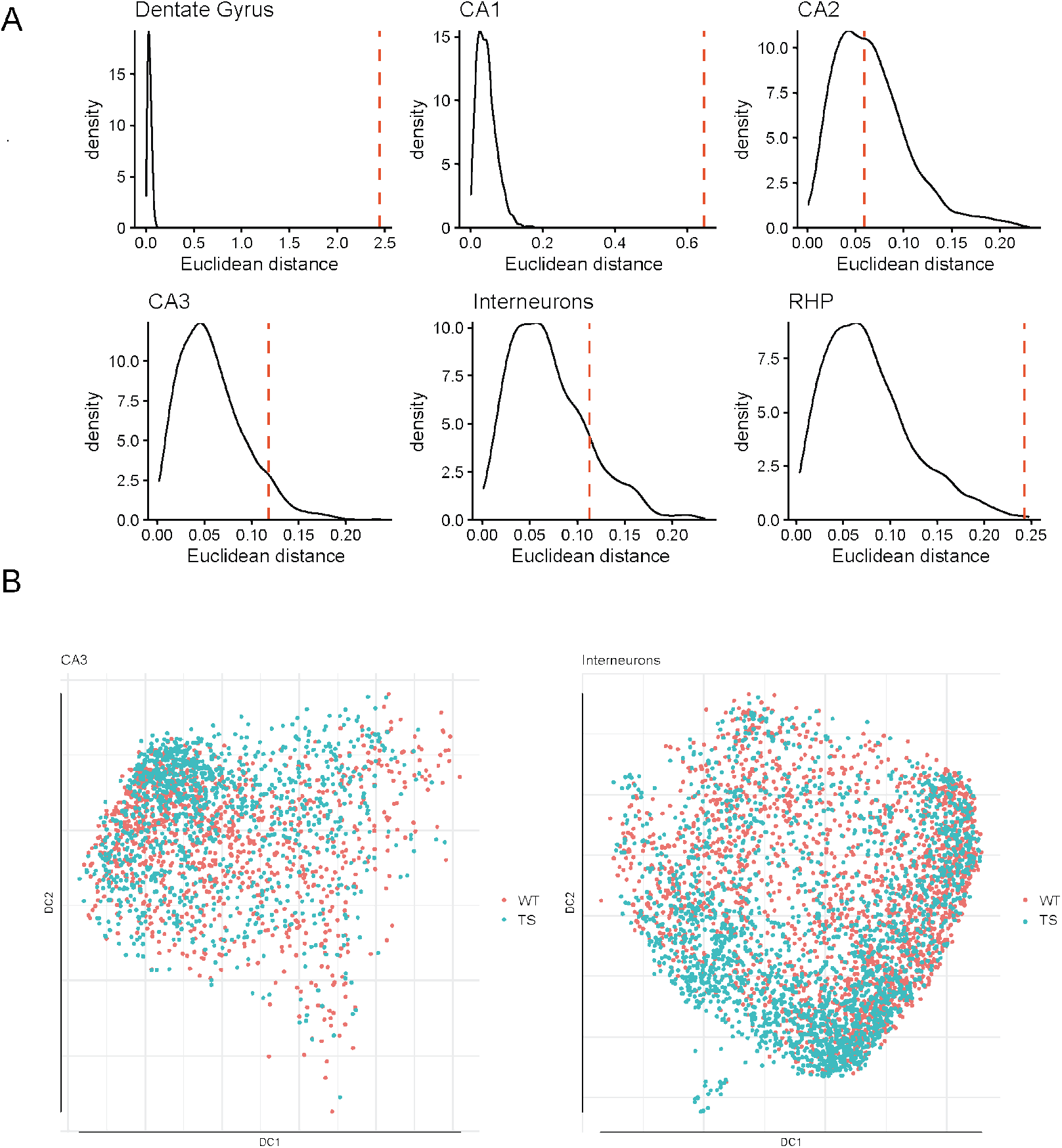
Dentate gyrus transcriptomic shift quantification. **(A)** Euclidean distance between WT and TS single cells for each cellular subtype. The density plot represents the null distribution of the distances between the single cells belonging to each cluster and the red line indicates the euclidean distance between the WT and TS centroid of each cluster. **(B)** Representation of the two-dimensional embedding of WT and trisomic cells by Diffusion map from CA3 pyramidal neurons (left) and interneurons (right).

**Supplementary Figure 6.**
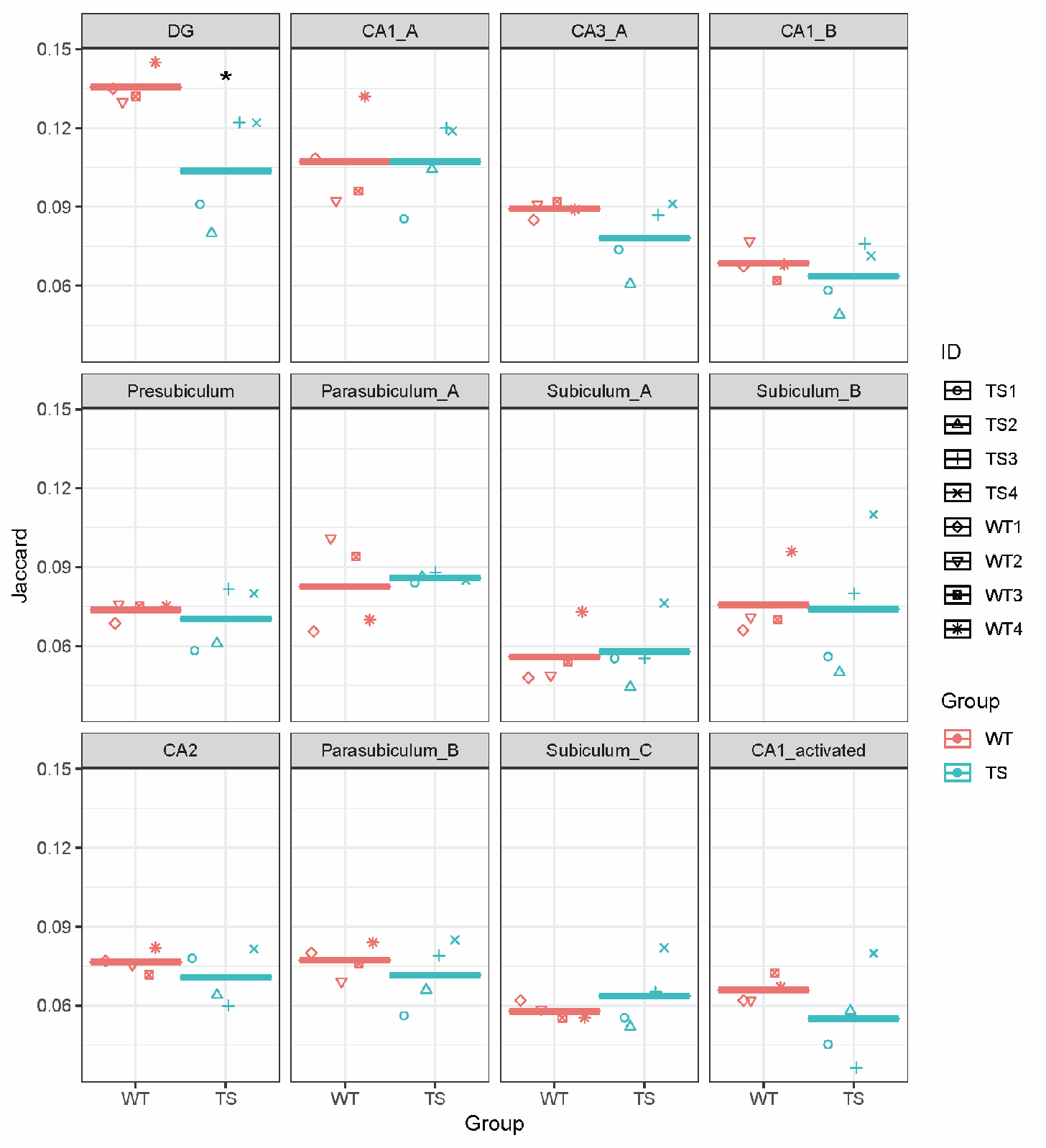
Jaccard values of the overlap between the cluster markers of each individual and the 300 most differentially expressed genes in each hippocampal subregion identified in the Allen Brain Atlas. * p < 0.05; Student t-test.

**Supplementary Figure 7.**
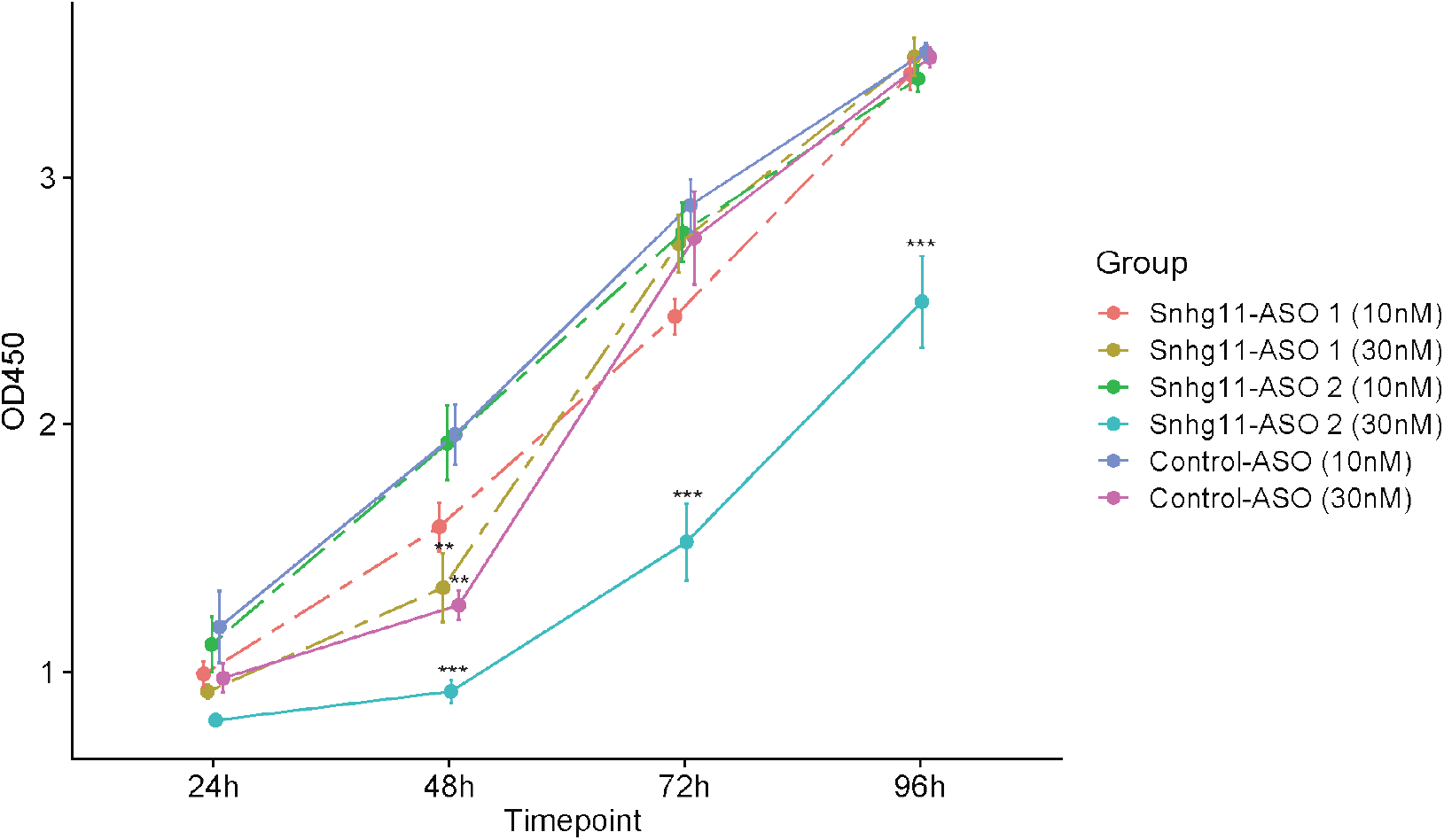
CCK8 quantification of cell proliferation for 96 hours after transfection of N2a cells with Control-ASO or with either ASO targeting Snhg11 (Snhg11-ASO 1 and Snhg11-ASO 2). * p< 0.05, ** p<0.01, *** p<0.001; TukeyHSD test.

**Supplementary Figure 8.**
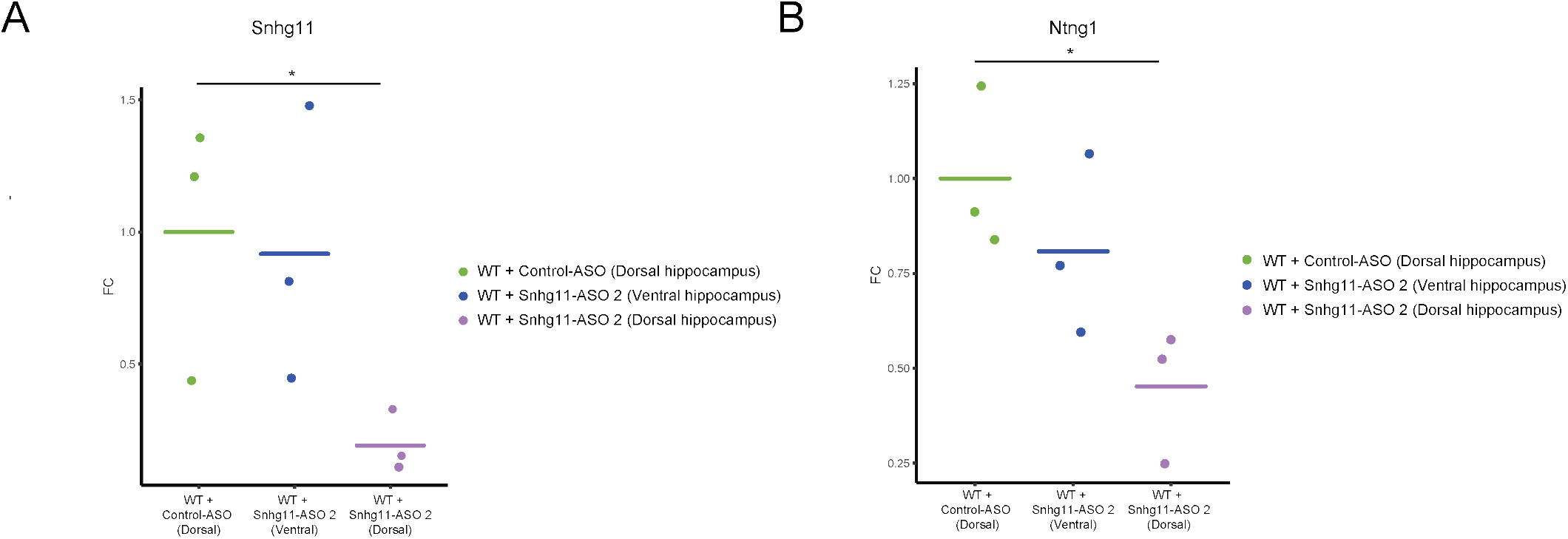
Gene expression quantification of Snhg11 and Ntng1 after ASO injection. **(A)** *RT-qPCR analysis of Snhg11 expression in the dorsal and ventral hippocampus of injected animals*. **(B)** *RT-qPCR analysis of Ntng1 expression in the dorsal and ventral hippocampus of injected animals. * p< 0*.*05*, Mann-Whitney-Wilcoxon test.

**Supplementary Figure 9.**
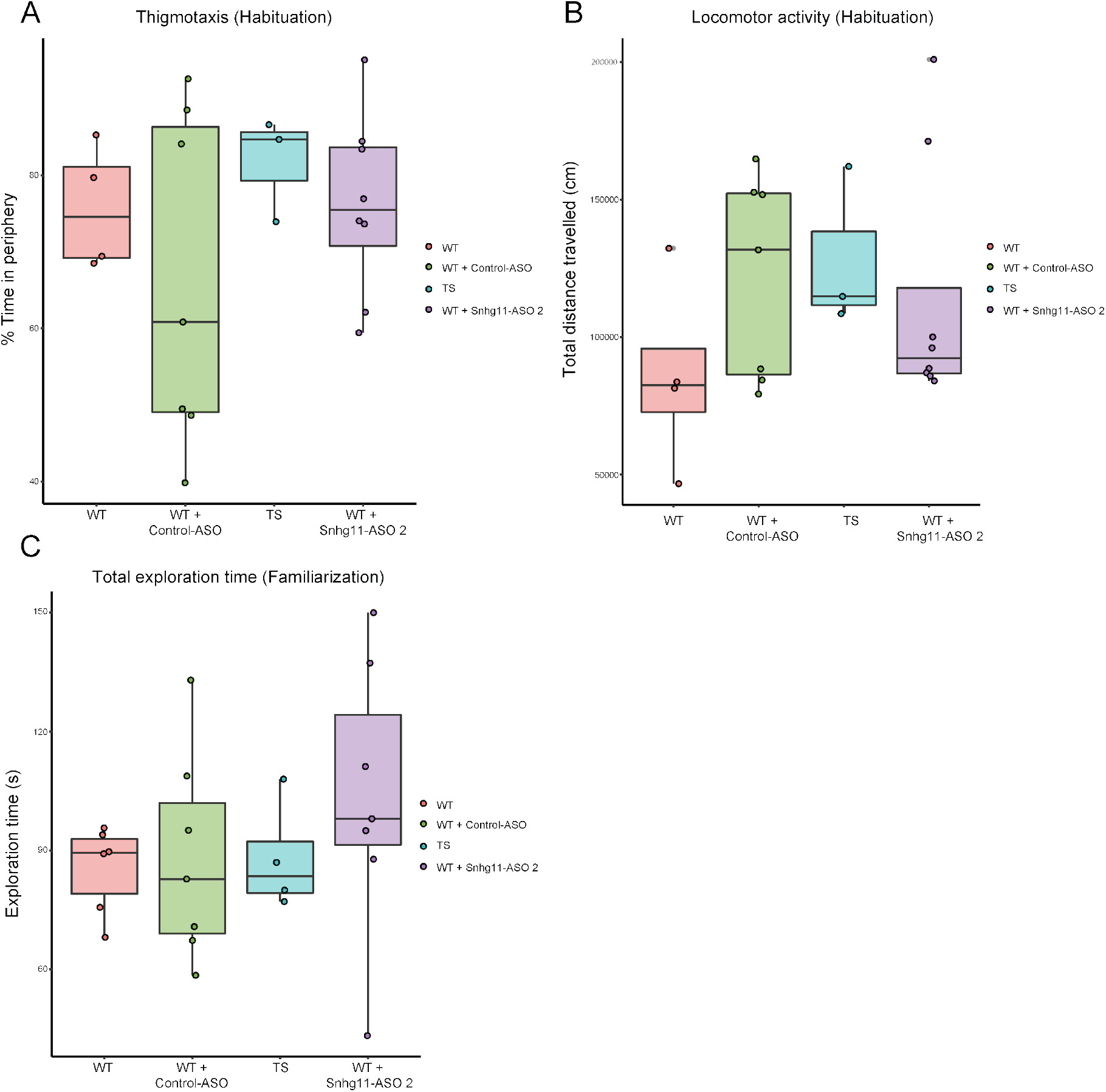
Novel object recognition performance in the habituation and familiarization sessions of NOR in WT, WT injected either with NC or ASO and TS mice. Boxplots depicting the experimental variables: **(A)** Percentage of time in periphery (Thigmotaxis) during habituation, **(B)** Total distance travelled in the habituation session and **(C)** Total exploration time recorded in the familiarization session.

